# High throughput droplet single-cell Genotyping of Transcriptomes (GoT) reveals the cell identity dependency of the impact of somatic mutations

**DOI:** 10.1101/444687

**Authors:** Anna S. Nam, Kyu-Tae Kim, Ronan Chaligne, Franco Izzo, Chelston Ang, Ghaith Abu-Zeinah, Nathaniel D. Omans, Justin Taylor, Alessandro Pastore, Alicia Alonso, Marisa Mariani, Juan R. Cubillos-Ruiz, Wayne Tam, Ronald Hoffman, Joseph M. Scandura, Raul Rabadan, Omar Abdel-Wahab, Peter Smibert, Dan A. Landau

**Affiliations:** Department of Pathology and Laboratory Medicine, Weill Cornell Medicine, New York, NY, USA; Division of Hematology and Medical Oncology, Department of Medicine and Meyer Cancer Center, Weill Cornell Medicine, New York, NY, USA; New York Genome Center, New York, NY, USA; Tri-Institutional Training Program in Computational Biology and Medicine, New York, NY, USA; Human Oncology and Pathogenesis Program, Memorial Sloan Kettering Cancer Center, New York, NY, USA; Division of Hematology and Medical Oncology, Department of Medicine, Epigenomics Core Facility, Weill Cornell Medicine, New York, NY, USA; Department of Obstetrics and Gynecology, Weill Cornell Medicine, New York, NY, USA; Division of Hematology/Medical Oncology, Department of Medicine, Tisch Cancer Institute, Icahn School of Medicine at Mount Sinai, New York, NY; Department of Systems Biology, Columbia University Medical Center, New York City, New York, USA; Technology Innovation Lab, New York Genome Center, NY, USA; Institute for Computational Biomedicine, Weill Cornell Medicine, New York, NY, USA

**Keywords:** Single-cell, RNA-seq, genotyping, hematopoiesis, myeloproliferative neoplasms, somatic evolution

## Abstract

Defining the transcriptomic identity of clonally related malignant cells is challenging in the absence of cell surface markers that distinguish cancer clones from one another or from admixed non-neoplastic cells. While single-cell methods have been devised to capture both the transcriptome and genotype, these methods are not compatible with droplet-based single-cell transcriptomics, limiting their throughput. To overcome this limitation, we present single-cell Genotyping of Transcriptomes (GoT), which integrates cDNA genotyping with high-throughput droplet-based single-cell RNA-seq. We further demonstrate that multiplexed GoT can interrogate multiple genotypes for distinguishing subclonal transcriptomic identity. We apply GoT to 26,039 CD34^+^ cells across six patients with myeloid neoplasms, in which the complex process of hematopoiesis is corrupted by *CALR*-mutated stem and progenitor cells. We define high-resolution maps of malignant versus normal hematopoietic progenitors, and show that while mutant cells are comingled with wildtype cells throughout the hematopoietic progenitor landscape, their frequency increases with differentiation. We identify the unfolded protein response as a predominant outcome of *CALR* mutations, with significant cell identity dependency. Furthermore, we identify that *CALR* mutations lead to NF-κB pathway upregulation specifically in uncommitted early stem cells. Collectively, GoT provides high-throughput linkage of single-cell genotypes with transcriptomes and reveals that the transcriptional output of somatic mutations is heavily dependent on the native cell identity.

## INTRODUCTION

Somatic mutations underlie the development of clonal outgrowth and malignant transformation^1-5^. Ongoing clonal evolution through acquisition of further genetic alterations often result in multiclonal cancer populations^6-10^. Transcriptional read-outs are critical for the study of the molecular basis of these processes. However, clonally-derived populations often lack cell surface markers that distinguish them from normal cells or that can help distinguish subclones, limiting the effectiveness of bulk RNA sequencing for investigating the clonal architecture of malignant cell populations. For example, myeloproliferative neoplasms (MPNs) have been shown to result from recurrent somatic mutations in *JAK2*, *CALR* and *MPL*, which are present across multiple progenitor classes, including CD34^+^, CD38^-^ hematopoietic stem progenitor cells (HSPC) and downstream progenitor cells including megakaryocytic-erythroid progenitors (MEP)^11,12^. These malignant clones often represent a subset of the bone marrow progenitor population without distinctive cell surface markers to distinguish them from non-neoplastic hematopoietic cells. Furthermore, MPNs are known to undergo clonal diversification with subclones harboring driver gene mutations^11,13,14^, without a tractable avenue for isolation of subclones based on cell surface markers. Therefore, our current understanding of the impact of the somatic mutations in MPN has relied primarily on cell lines, genetic mouse models, and profiling of expanded patient-derived cells that may undergo significant alteration during *in vitro* expansion.

Recently, advanced methods have been developed to capture both the transcriptional information and genotype at the single-cell level. Simultaneous single-cell transcriptomic (scRNA-seq) analysis and *BCR-ABL* detection was performed using *BCR-ABL*-specific primers during reverse transcription and amplification steps using the Smart-seq2 platform^15^. In another study, single-cell targeted genotyping of *EGFR* combined with gene expression and DNA methylation analyses was performed in lung adenocarcinoma cells by adapting the Fluidigm platform and amplifying DNA at the locus of interest^16^. While these methods provide rich annotations of single-cells, they are limited in throughput, providing information in the order of hundreds of cells. The lower throughput of these methods is especially limiting when profiling highly complex cellular populations, such as hematopoietic progenitors, which consist of a large number of distinct cellular subsets^17-20^.

Droplet-based sequencing has been shown to be a powerful approach, driving library preparation and sequencing costs down on a per-cell basis and providing high resolution maps of complex population structures through the single-cell transcriptomic profiling of thousands of cells^21-23^. Nevertheless, the most commonly used droplet-based methods determine transcript abundance by counting short sequence tags towards the 5’ or 3’ end of the transcript, limiting the ability to link genotype information with single-cell transcriptional identities.

To overcome this limitation, we developed single-cell Genotyping of Transcriptomes (GoT), as well as its analytic pipeline IronThrone GoT, to link genotypes of expressed genes to transcriptional profiling of thousands of single-cells, leveraging the throughput of droplet-based sequencing. We performed gene-specific enrichment from full-length barcoded cDNAs generated at intermediate steps of common droplet-based single-cell methods^22,24^. We applied GoT to MPNs which result from acquisition of recurrent somatic mutations in the HSPCs. By profiling 26,039 CD34^+^ cells from six patients, we superimpose the genotype information on the map of highly complex and heterogeneous pool of hematopoietic progenitors. GoT revealed that *CALR* mutations resulted in increased proliferation in committed progenitors, especially megakaryocytic progenitors (MkPs), associated with increased mutated cell fraction. In uncommitted HSPCs, the NF-κB pathway was upregulated, serving as a potential mechanism for increased self-renewal. We found that *CALR* mutations resulted in variable upregulation of IRE1-mediated unfolded protein response, dependent on cell identity. These data revealed that MPN mutations in hematopoietic progenitor cells do not lead to uniform transcriptional output, but rather show strong dependence on progenitor subset identity.

## RESULTS

### Single-cell Genotyping of Transcriptomes (GoT) provides coupling of targeted cDNA genotyping with single-cell whole transcriptomes

Gene expression leads to a significant natural information enrichment, as only a small fraction of the genome is transcribed. Moreover, pan-cancer analysis revealed that for most driver mutations the expressed mRNA variant allele fraction (VAF) is equal or higher compared to genomic DNA VAF (**Supplementary Fig. 1**). Thus, mRNA-based detection may empower the genotyping of transcribed genes in single cells, overcoming the inherent sparsity of single-cell genomics. However, high-throughput digital scRNA-seq methods, such as Drop-seq or the 10x Genomics Chromium 3’ platform, are by design heavily 3’ biased, providing the sequence information of only a short fragment, typically in the last exon.

To link genotypes to single-cell transcriptomes in high throughput droplet-based platforms, we devised a strategy to pair targeted genotyping with single-cell whole transcriptomics (GoT). First, we add gene-specific primers during the cDNA amplification step of the 10x Chromium procedure to promote amplification of even lowly-expressed genes of interest (**Fig. 1a**). Second, after generation of the cDNA library, a small portion of the cDNA (~10%) is aliquoted for targeted genotyping. Locus specific primers are designed based on known somatic mutations identified from bulk DNA genotyping of the sample, and used to amplify the locus of interest together with the generic forward SI-PCR primer (10x Genomics) to retain the cell barcode (CB) and unique molecule identifier (UMI; **Fig. 1a**). The targeted amplicon library is subsequently spiked back into the 10x gene expression library to be sequenced together, or may be alternatively sequenced separately. Finally, we interrogate target amplicon reads for mutation status at the locus of interest, and link the genotype information to single-cell gene expression profiles via shared cell barcode information (see online methods, **Supplementary Fig. 2a,b**).

**Figure 1.**
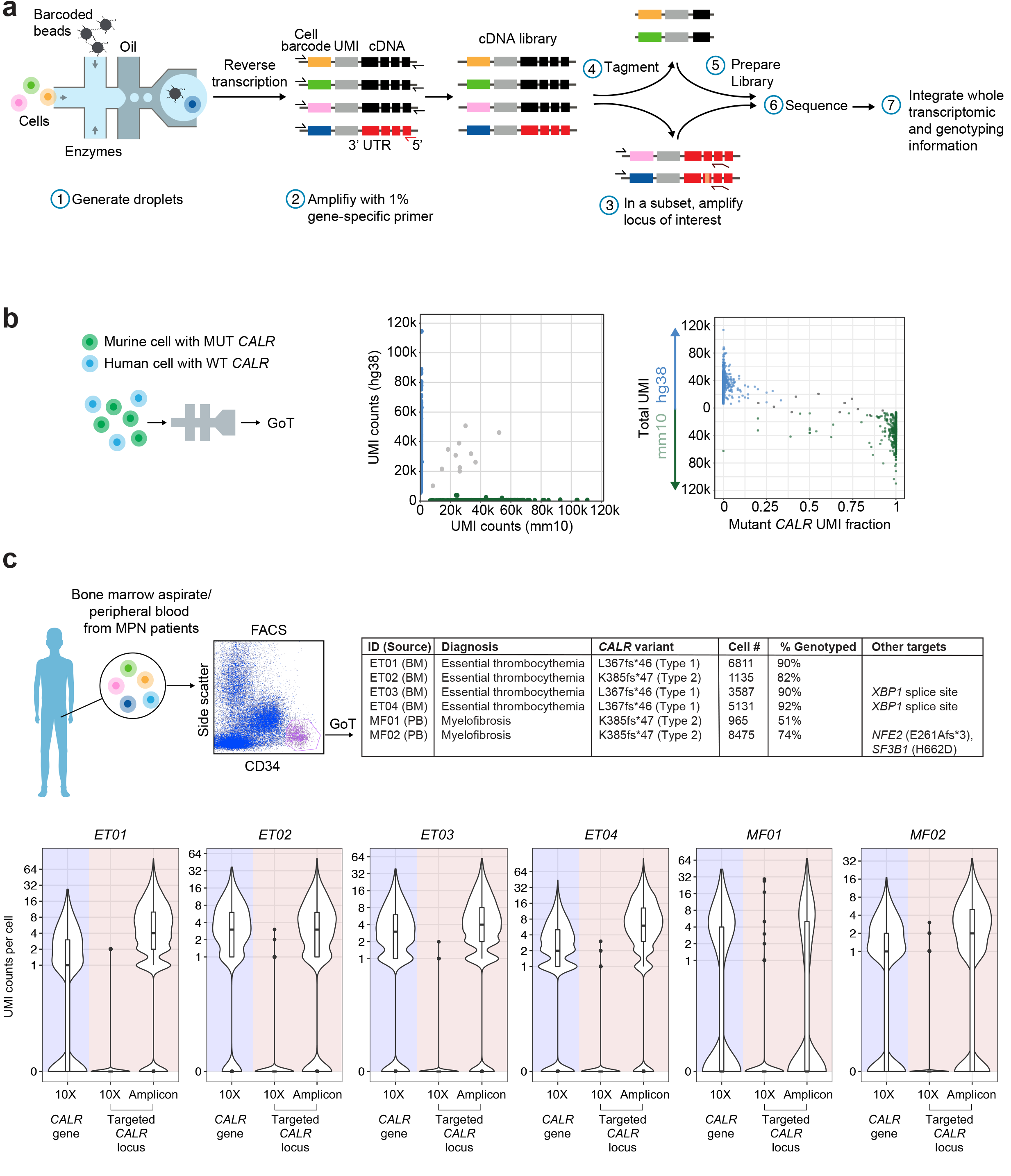
Single-cell Genotyping of Transcriptomes (GoT) provides accurate coupling of targeted cDNA genotyping with single-cell whole transcriptomes. **(a)** Schematic representation of GoT. Standard 10x procedure (step 1) is carried out until the cDNA amplification step in which we add gene-specific primers (step 2). After generation of the cDNA library, a small portion of the cDNA (~10%) is aliquoted for targeted genotyping. Locus-specific primers are used to amplify the locus of interest together with generic forward SI-PCR primers to retain the cell barcode (CB) and unique molecule identifier (UMI) (step 3). The remainder of the library undergoes tagmentation (step 4). After Illumina compatible libraries are prepared (step 5), the targeted amplicon library is sequenced in tandem with the standard 10x library or separately (step 6). **(b)** Species-mixing experiment in which mouse cells with a human mutant *CALR* transgene were mixed with human cells with a human wildtype *CALR* transgene. Cells identified as human or murine based on scRNA-seq reads that mapped to either the human or mouse genome (left). Murine (green) vs. human (blue) genome alignment of 10x data (y-axis) with genotyping data by GoT (x-axis, right). Multiplet cells are shown in gray. **(c)** Summary of GoT data from FACS-sorted CD34^+^ bone marrow aspirate/peripheral blood cells from patients with myeloproliferative neoplasms (MPN, top). Number of UMIs per cell of *CALR* gene from 10x data (left) or targeted *CALR* locus from 10x (middle) or GoT (right) from each of the ET patient sample (bottom). BM, bone marrow; PB, peripheral blood.

To test the ability of GoT to accurately read-out joint single-cell genotypes and transcriptomes, we performed a species-mixing experiment, whereby mouse cells (Ba/F3) with a human mutant *CALR* (type 1, L367fs*46) transgene were mixed with human cells (UT-7) with a human wildtype *CALR* transgene (**Fig. 1b**)^25^. Consistent with precision genotyping, the vast majority of cell barcodes with transcripts mapped to the mouse genome were enriched for mutant *CALR*, whereas cell barcodes with scRNA-seq reads annotated to the human genome were enriched for wildtype *CALR*. A mean (± standard deviation) of 83.6 (± 95.3) *CALR* UMIs were detected per cell in the amplicon data, with 52 (± 16.3) reads per UMI.

The species mixing data provided an opportunity to optimize the analytical assignment of genotypes to cells, to overcome technical sources of noise such as PCR errors and PCR recombination, which often accompany targeted amplification. We integrated targeted amplicon measures including base quality, number of mismatches, CB matching, and number of duplicate reads per UMI, to determine optimized parameters that maximize the number of genotyped cells while minimizing genotype mis-assignment (see online methods, **Supplementary Fig. 3a-e**). By applying these optimized parameters to the species-mixing experiment data, we demonstrate that GoT provides genotyping information for 97.5% of cells, with correct genotyping (i.e., matching the expected species) in 96.7% (**Fig. 1b**).

### GoT provides joint genotype and whole transcriptome information for thousands of CD34^+^ cells from primary patient samples

Human CD34^+^ HSPCs and closely-related lineage committed progenitor cells harbor significant cell type complexity, with multiple distinct progenitor populations that trace the differentiation from early HSCs to mature blood cells. However, the lack of FACS-sortable surface markers that effectively distinguish wildtype from mutant cells in MPN prevents integration of this complexity with genotypic information.

We therefore applied GoT to FACS-sorted viable CD34^+^ cells from six patients with *CALR*- mutated MPNs, including cryopreserved bone marrow aspirate samples from four patients with essential thrombocythemia (ET). These four patients were not treated with disease-modifying therapy at the time of biopsy (see clinical information in **Supplementary Table 1**). ET manifests with increased circulating platelet counts, known to result from megakaryocytic proliferation, and is associated with heterozygous *CALR* mutations in 25% of patients^26^. Disease-causing *CALR* mutations are frameshifts that result in a novel C-terminus which is thought to confer an activating interaction with the thrombopoietin receptor MPL, thus resulting in megakaryocytic proliferation^25, 27-30^. However, the role of *CALR* mutations in perturbing the early HSPC differentiation in patients is still largely unknown. Thus, a total of 16,664 cells from the four ET patients were sequenced with GoT and included in downstream analysis after quality filters (online methods).

Of the 16,664 cells, 15,001 cells (90.0%) were linked to targeted *CALR* amplicon data, compared to 137 cells (0.8%) by interrogation of the *CALR* sequence in the conventional 10x data (**Fig. 1c**, **Supplementary Fig. 4**). The *CALR* locus of interest was covered by a median (± median absolute deviation) of 5 (± 4.4) targeted amplicon UMIs per cell. Across the four patient samples, mutant progenitor cells accounted for 26.2%-68.8% of the cells, with good agreement with matched bulk *CALR* exon sequencing (e.g., ET01 VAF of 15% for the heterozygous *CALR* mutation and 26.2% *CALR* mutant cells in GoT). Nevertheless, we note that cells heterozygous for mutations may be mis-assigned as wildtype due to the partial sampling of the mRNA pool in droplet scRNA-seq or due to transcriptional bursts^31^. To overcome these potential biases, in our analyses, we undertook down-sampling, as well as systematic examination of the relationship of genotyped-based findings to varying the minimum UMI number thresholds (**Supplementary Fig. 5a-c**). We further note that in order not to compromise the scRNA-seq data, the 10x cDNA library underwent an extra cycle of PCR beyond the manufacturer’s recommended number of cycles before subtracting ~10% of the cDNA library for GoT. Indeed, the GoT procedure did not result in significant loss of genes or UMIs per cell in comparison to published data of CD34^+^ selected cells from the standard 10x library **(Supplementary Fig. 6a-c)**^24^.

### GoT maps the hematopoietic differentiation of *CALR* mutant vs. wildtype progenitor cells within the same individuals

To interrogate the cellular identities of hematopoietic progenitors in these ET samples, we performed clustering agnostic to the genotyping information, based on the transcriptome information alone, using the Seurat package^32^. We identified 9-19 distinct clusters in the four samples, which were annotated according to marker genes identified by Velten *et al* (2017)^17^ (representative t-distributed stochastic neighbor embedding (t-SNE) from patient ET01 in **Fig. 2a,** and clustering heatmap in **Supplementary Fig. 7a)**. Our data are consistent with previous reports of normal donor bone marrow CD34^+^ progenitor cell clustering, and cluster identities were readily assigned based on progenitor identity markers identified in previous scRNA-seq reports.

**Figure 2.**
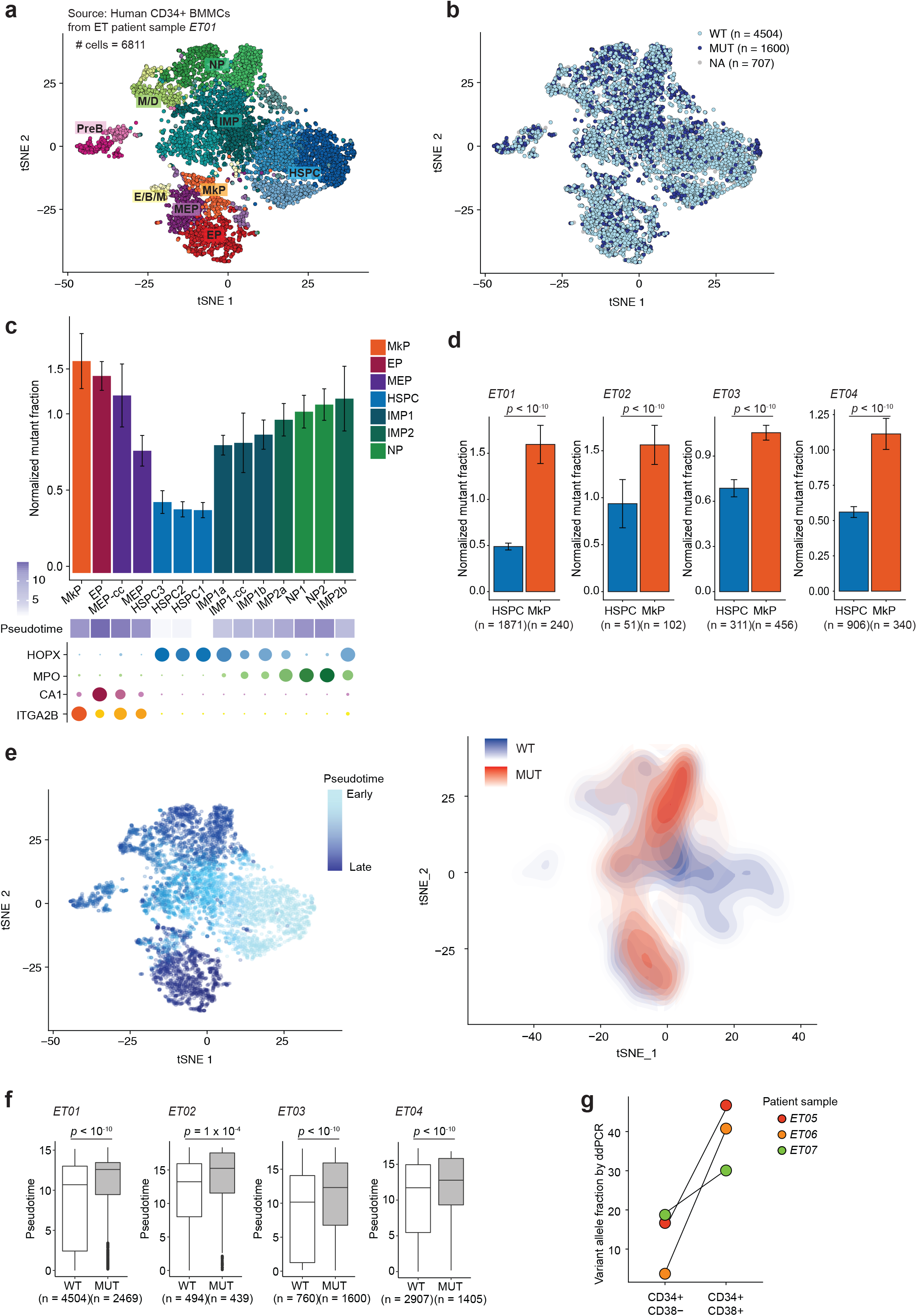
GoT reveals higher frequency mutated cells in committed progenitor subsets compared to HSPCs. **(a)** tSNE projection of the CD34^+^ bone marrow mononuclear cells (BMMCs) from patient with essential thrombocythemia (ET01) using the Seurat package (see online methods)^32^. **(b)** tSNE projection of the CD34^+^ cells from patient ET01 showing genotyping information based on GoT data. Cells without any GoT data are labeled NA (not available). **(c)** Frequency of mutant cells within each progenitor cluster from sample ET01 after down-sampling targeted amplicon data to one *CALR* UMI per cell and normalizing by total mutant frequency across the entire dataset. Mean ± standard deviation (SD) of 100 down-sampling iterations. Progenitor clusters are annotated by average pseudotime, and by expression of marker genes (bottom, size represents percentage of expressing cells; color intensity represents average expression level). **(d)** Normalized frequency of mutant cells in hematopoietic stem progenitor cell (HSPC) and megakaryocytic progenitor (MkP) clusters. Mean ± standard deviation (SD) after 100 down-sampling iterations (Wilcoxon rank-sum test). **(e)** tSNE projection of the CD34^+^ cells from patient ET01 showing pseudotime assignment (left) and density plot of mutant and wildtype cells (right). **(f)** Pseudotime comparison between wildtype (WT) and mutant (MUT) cells. *P*-values from generalized linear model including mutation status and total number of amplicon UMIs per cell. **(g)** Variant allele fraction in CD34^+^, CD38^-^ and CD34^+^, CD38^+^ FACS-sorted peripheral blood cells from patients with ET determined by droplet digital PCR (ddPCR). HSPC, hematopoietic stem progenitor cells; IMP, immature myeloid progenitors; NP, neutrophil progenitors; M/D, monocyte-dendritic cell progenitors; E/B/M, eosinophil, basophil, mast cell progenitors; MEP, megakaryocytic-erythroid progenitors; MkP, megakaryocytic progenitors; EP, erythroid progenitors; PreB, pre-B cells; WT, wildtype; MUT, mutant.

Next, we projected the genotype information onto the progenitor cluster maps. We observed that in all four ET affected individuals, mutated cells were observed across all the CD34^+^ stem and progenitor clusters, consistent with previous PCR analysis of *CALR* in FACS-sorted CD34^+^ subsets (**Fig. 2b** and **Supplementary Fig. 7b**)^11^. Notably, the presence of cells mutated for *CALR* did not result in novel independent clusters. These data confirmed that scRNA-seq alone is insufficient to distinguish mutant from wildtype cells, demonstrating the need for genotyping information. Furthermore, GoT demonstrated that *CALR* mutations in ET patients impact the entire hematopoietic differentiation hierarchy, without forming new distinct cellular identities.

While mutant cells were observed across all progenitor clusters, we hypothesized that the frequency of mutated cells may vary between clusters. Notably, as *CALR* expression is higher in committed progenitor clusters, we applied both amplicon UMI down-sampling and UMI threshold sensitivity analysis to exclude sampling biases as a potential confounder leading to underestimation of mutated cell frequency in the lowly *CALR* expressing HSPCs (**Fig. 2c,d** and **Supplementary Fig. 5c**). Interestingly, the fraction of *CALR* mutated cells was higher in committed progenitors, in particular MkPs, compared to HSPC clusters across all patient samples (each *P*-value <10^-10^, **Fig. 2d**). To further explore this observation, independent of cluster assignment, we performed a differentiation pseudo-temporal ordering (pseudotime) analysis using Monocle^33^, and found that *CALR* mutated cells were enriched in cells with later pseudotime points compared to wildtype cells (**Fig. 2e**). Consistent with this finding, mutant cells were on average associated with later pseudotime points compared with wildtype *CALR* cells (*P*-values < 10^-4^, including in multi-variable model that explicitly accounts for the number of genotyping amplicon UMI per cell, **Fig. 2f** and **Supplementary Fig. 5a**). This observation was orthogonally validated via bulk genomic DNA VAF analysis of *CALR* mutation in FACS-sorted CD34^+^, CD38^-^ vs. CD34^+^, CD38^+^ progenitor subsets from additional ET patients via droplet digital PCR, showing a lower frequency of *CALR* mutations in HSPCs compared to that of CD34^+^, CD38^+^ progenitors (*P*-value 0.02, **Fig. 2g**). This finding suggested that while *CALR* mutations likely arise in uncommitted HSCs and, therefore, propagate to populate the entire differentiation tree, the impact of *CALR* mutations on fitness increases with differentiation.

### GoT transcriptional comparison of cluster-matched *CALR* mutant vs. wildtype cells reveals differentially increased proliferation in mutant progenitor subsets

Due to the high-resolution mapping of cellular subtypes coupled with genotyping, GoT uniquely enables direct transcriptional program comparison between mutant and wildtype cells, not only within the same sample, but also *within the same progenitor cluster*. For example, single-cell cell cycle-related gene signature may provide an accurate estimate of the proportion of cells undergoing cell replication^22,34^. Consistent with the above suggestion of increased fitness of *CALR* mutations in MkPs compared with HSPCs, *CALR* mutant HSPCs exhibited only a modest increase in single-cell cell cycle gene expression^35^ compared to wildtype HSPCs (*P*-value 0.007, mean fold change 1.3, 95% confidence interval, 1.1-1.5; **Fig. 3a, Supplementary Table 2**). In contrast, mutant cells in the MkP cluster exhibited a greater increase in the cell cycle-related gene expression compared with their wildtype counterparts (*P*-value 10^-5^, fold change 3.1 [2.1- 4.0]; **Fig. 3a**). While increased cell cycle score in MkPs was most pronounced in ET02, ET03 and ET04, mutant MkPs in sample ET01 did not display a significantly higher proliferative status than wildtype MkPs (**Supplementary Fig. 8a**). Notably, this patient showed a lower platelet count in the peripheral blood—just above the upper limit of normal—compared to the other three patients (519K/μL compared with 1060K/μL, 1252K/μL and 650K/ μL). These data suggest a correlation between MkP mutant vs. wildtype differential cell cycle score and the platelet count (**Fig. 3b**). Thus, interrogation of the early progenitor mutant and wildtype cells may correlate with clinical phenotypes and inform our understanding of patient-to-patient variability despite shared mutated genotypes.

**Figure 3.**
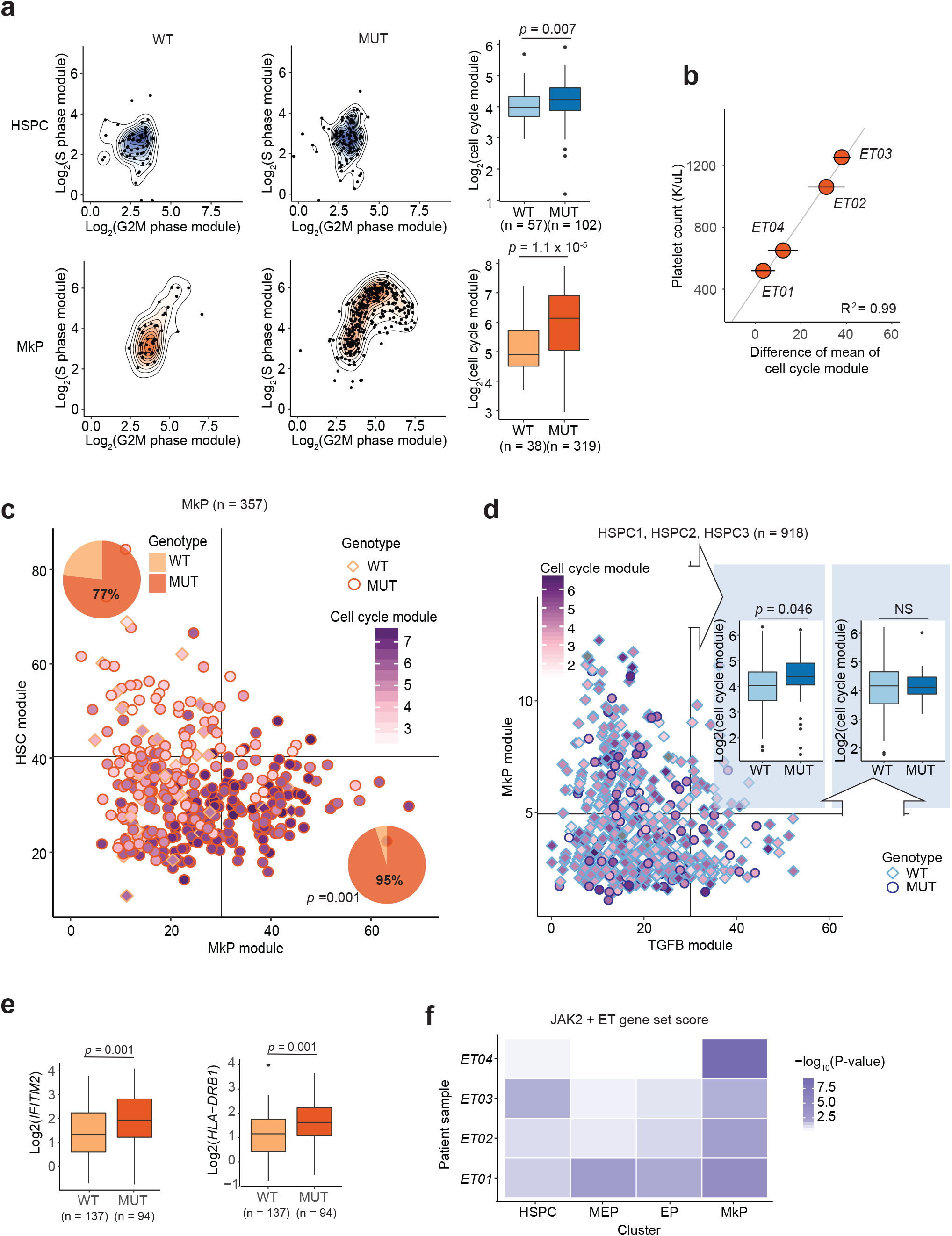
GoT reveals higher proliferative impact of *CALR* mutation in megakaryocytic progenitor subset compared to uncommitted HSPCs. **(a)** S phase and G2M phase gene module expression in wildtype (WT) vs. mutant (MUT) cells in HSPC and MkP clusters from representative patient sample (ET03). Cell cycle module score represents sum of S phase and G2M phase gene module expression (Wilcoxon rank-sum). **(b)** Correlation between the difference of mean cell cycle score (± standard error) between WT and MUT MkP cells, and respective patients’ platelet count at the time of bone marrow biopsy. **(c)** MkP WT and MUT cells from a representative sample (ET03) mapped based on MkP gene module and HSC gene module expression, annotated with cell cycle score. Pie charts show frequency of WT vs. MUT cells in HSC^lo^MkP^hi^ and HSC^hi^MkP^lo^ populations (Fisher’s exact test). **(d)** WT and MUT cells from HSPC1, HSPC2, and HSPC3 clusters from patient with the highest HSPC cluster complexity (ET01) mapped based on MkP gene module and TGFβ signaling pathway module, annotated with cell cycle score. Cell cycle module score comparison between WT and MUT HSPCs in MkP^hi^TGFβ^lo^ and MkP^lo^TGFβ^hi^ populations (Wilcoxon rank-sum test). **(e)** Expression of example genes upregulated in *JAK2*-mutated essential thrombocythemia (ET) patient-derived cultured cells in *CALR* WT and MUT MkP cells (Wilcoxon rank-sum test). **(f)** Heatmap of - log10(*P-*value) (Wilcoxon rank-sum test) of expression level of genes that are upregulated in *JAK2*-mutated ET patient-derived cultured cells, between WT and MUT cells in progenitor clusters.

In addition, cell-to-cell variation may exist even within progenitor subsets. For example, we observed that MkPs represent a heterogeneous population composed of cells that are in transition from less differentiated cells with higher expression of HSC genes^36^ (HSC^hi^MkP^lo^), to more committed MkP cells displaying high expression of MkP-specific genes (HSC^lo^MkP^hi^; **Fig. 3c**, **Supplementary Table 2**). Notably, even within the MkP cluster, HSC^lo^MkP^hi^ cells showed increased cell cycle gene expression, and had a higher proportion of mutant cells compared to the HSC^hi^MkP^lo^ group (*P*-value 0.001, Fisher’s exact test). Similarly, we interrogated uncommitted HSPC clusters for the expression of MkP genes, indicating megakaryocyte/platelet-primed HSPCs, vs. genes in the TGFβ signaling pathway (KEGG), which maintains HSC quiescence^37^ (**Fig. 3d**). Mutant cells within the platelet-primed HSPCs (MkP^hi^TGFβ^lo^) expressed higher cell cycle-related genes than the wildtype MkP^hi^TGFβ^lo^ HSPCs, whereas the mutant and wildtype HSPCs in a more quiescent state (MkP^lo^TGFβ^hi^) showed no difference in cell cycle score. This finding further emphasized that downstream output of *CALR* mutation shows a strong dependency on cell state and imparts a greater proliferative advantage in cells that are more differentiated. These data also suggested that *CALR* mutations skews differentiation toward megakaryocytes early in hematopoiesis.

### *CALR* mutation imparts differential transcriptional output as a function of cellular identity

In addition to the cell cycle transcriptional signature, we tested whether *CALR*-mutated hematopoietic progenitors displayed increased expression of genes that were upregulated in single-cell clones of cultured CD34^+^ cells from patients with *JAK2*-mutated ET^38^. We observed an increase in expression of genes in this gene set, including those of the interferon pathway (e.g. *IFITM2, HLA-DRB1*; **Fig. 3e**). Notably, the gene set increase was most significant in MkP clusters (combined *P*-value <10^-10^, Fisher’s method) compared to the other subsets (**Fig. 3f**). This result supports previous reports suggesting that *JAK2* and *CALR* mutations partially converge through activation of similar downstream pathways^39^.

GoT data further offers a unique opportunity for *de novo* differential gene expression discovery, by examining gene expression in wildtype vs. mutant cells within the same progenitor subset. Crucially, in this analysis, the wildtype cells serve as an ideal comparison set, as they share all potential environmental and patient-specific variables as the mutated cells. To identify gene expression changes associated with *CALR* mutations, we performed a differential expression analysis comparing progenitor-cluster matched mutant and wildtype cells.

We identified 37 genes to be differentially expressed between mutant and wildtype MkPs (FDR <0.25; **Fig. 4a**). Mutated MkPs upregulate *HSPA5* that encodes BiP, a key player in protein quality control that modulates the activities of the three transmembrane transducers of the unfolded protein response (UPR): PERK, IRE1, and ATF6^40^. Consistently, *XBP1*, another chief regulator of the UPR, was also upregulated in MkP (**Fig. 4a**)^40^. Both *HSPA5* and *XBP1* expression are upregulated by ATF6 activation, and indeed, an enrichment analysis of genes upregulated in *CALR-*mutant MkPs showed enrichment of ATF6-mediated UPR (**Fig. 4a,b**). UPR in this context may signal ER stress in response to misfolded proteins, as the *CALR* chaperone activity may be compromised^41^.

**Figure 4.**
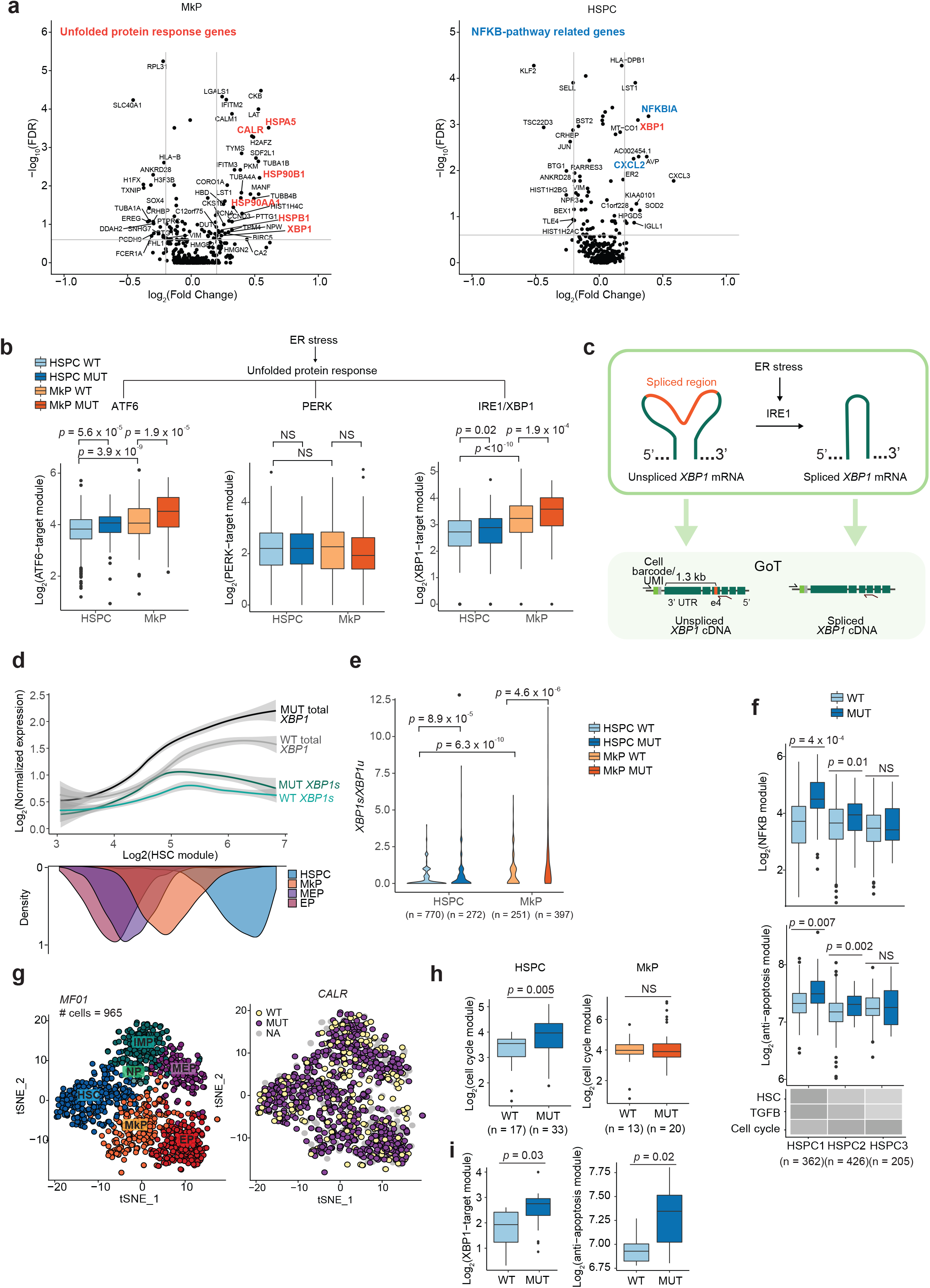
Downstream effects of *CALR* mutation is cell identity dependent. **(a)** Volcano plots of differentially expressed genes in megakaryocytic progenitors (MkP, left) and hematopoietic stem progenitor cells (HSPC, right), common across patient samples ET01, ET02, ET03, and ET04. *P*-values were combined using Fisher combine test and multiple hypothesis correction was performed (Benjamini-Hochberg FDR). Average of log_2_(fold change) across samples is shown. **(b)** Aggregate expression of ATF6-, PERK- and XBP1-target genes in the unfolded protein response in WT and MUT HSPC and MkP cells from a representative patient sample (ET04). **(c)** Schematic representation of GoT applied to the unconventional splice site of *XBP1* mRNA by IRE1 in response to ER stress. **(d)** Local regression of total and spliced *XPB1* (*XBP1s*) expression in progenitor cells ordered by expression of HSC module (x-axis) from patient samples ET03 and ET04 where excess material allowed *XBP1* GoT. **(e)** Spliced *XBP1* to unspliced *XBP1* ratio (*XBP1s* UMI*/ XBPu* UMI+1) in WT and MUT HSPCs and MkPs from patient samples ET03 and ET04. **(f)** Aggregate expression of NF-κB signaling pathway-related genes and anti-apoptosis genes in WT and MUT cells from HSPC1, HSPC2, and HSPC3 subclusters (Wilcoxon rank-sum test). Heatmap shows normalized expression level of gene modules of all cells within each subcluster. **(g)** tSNE projection of the CD34^+^ peripheral blood mononuclear cells from patient with myelofibrosis (MF01) using the Seurat package (see online methods)^32^ showing progenitor identity assignment (left) and genotyping information (right) based on GoT data. Cells without any GoT data are labeled NA (not available). **(h)** Cell cycle module score comparison between WT and MUT HSPCs (left) and MkPs (right) from patient with myelofibrosis (MF01). **(i)** XBP1-target genes in UPR module scores (left) and anti-apoptosis module scores (right) in WT and MUT HSPCs with high expression (i.e. greater than the median value) of HSPC1-module (Wilcoxon rank-sum test). HSPC, hematopoietic stem progenitor cells; IMP, immature myeloid progenitors; NP, neutrophil progenitors; MEP, megakaryocytic-erythroid progenitors; MkP, megakaryocytic progenitors; EP, erythroid progenitors; WT, wildtype; MUT, mutant.

Notably, *XBP1* was also upregulated in differential expression analysis of mutant vs. wildtype HSPCs suggesting that UPR activation by *CALR* mutations extends to uncommitted progenitors (**Fig. 4a**). Consistently, ATF6-target genes were also increased in mutant HSPCs compared to wildtype HSPCs (**Fig. 4b**, **Supplementary Fig. 8b, Supplementary Table 2**)^42^.

UPR was previously demonstrated to be preferentially mediated in HSPCs through PERK resulting in enhanced apoptosis upon ER stress^43^. In contrast, committed progenitors have a robust IRE1 activity, correlated with survival in ER stress challenge^43^. Thus, we interrogated these branches of the UPR in *CALR* mutation-associated ER stress, and observed that while the HSPCs showed a trend of expressing higher levels of PERK pathway-related genes compared to the MkPs, the PERK-branch of the UPR was not significantly enhanced by mutated *CALR*- associated ER stress (**Fig. 4b, Supplementary Fig. 8b, Supplementary Table 2**)^43^. As IRE1 catalyzes the unconventional splicing of *XBP1* mRNA (*XBP1u*) into the active spliced form (*XBP1s*; **Fig. 4c**)^44^, to determine whether the IRE1-branch of the UPR is activated by *CALR* mutations, we repurposed GoT to probe for the spliced region of *XBP1* in CD34^+^ cells. Consistent with previous report^43^, GoT analysis of the *XBP1* spliced region demonstrated that while the total *XBP1* UMI was upregulated in HSPCs compared to committed progenitors, *XBP1s*/*XBP1u* ratio in HSPCs was significantly lower in HSPCs (**Fig. 4d,e**). *CALR* mutations robustly augmented the amount of *XBP1s* in MkPs (**Fig. 4d,e**). Remarkably, *CALR* mutations also resulted in enhanced *XBP1s*/*XBP1u* ratio in HSPCs, indicating increased IRE1 activity (**Fig. 4d,e**). Consistent with this finding, targets of XBP1 were upregulated in both mutant MkPs and HSPCs (**Fig. 4b, Supplementary Fig. 8b, Supplementary Table 2**)^45,46^. These data thus suggest that in the *CALR*-related UPR, IRE1 is activated in HSPCs, skewing the ER stress challenged stem cells toward survival.

### NF-κB pathway is upregulated in *CALR* mutated HSPCs

Differentially expressed gene analysis in the HSPCs also revealed upregulation of the NF-κB pathway in the HSPCs (*P*-value 0.001, **Supplementary Fig. 8c**), including upregulation of *CXCL2* and *NFKBIA* (**Fig. 4a**). We further analyzed the NF-κB gene set enrichment (**Supplementary Table 2**) in HSPC subclusters, which showed the strongest upregulation in mutant vs. wildtype HSPCs enriched for uncommitted HSCs, based on high expression of HSC-related and TGFβ pathway genes on the one hand, and low expression of cell cycle-associated genes on the other (**Fig. 4f**). The mutant cells in this subcluster also upregulated anti-apoptotic related genes (**Fig. 4f**, **Supplementary Table 2**). In contrast, NF-κB pathway genes were not upregulated in mutant cells in more differentiated progenitor subsets like MkPs (mean log2(expression) of 3.8 (3.7-3.9) in wildtype cells vs. 3.9 (3.8-4) in mutant cells, *P-*value 0.35), suggesting an HSPC-specific effect. NF-κB pathway activation has been previously associated with HSC self-renewal^47^, and thus, our data points at another potential mechanism linking *CALR* mutation and the outgrowth of HSCs.

Collectively, these findings demonstrate that *CALR* mutations have distinct and specific impact on the transcriptomic output depending on the transcriptional programs already in place that define cell identity (i.e., differential programs activated in committed progenitors vs. uncommitted HSPCs). Thus, it is imperative not only to examine the role of somatic mutations directly in patient samples, but also to analyze the downstream effect of these mutations in the context of cell identity.

As a proportion of *CALR*-mutated ETs eventually progresses to post-ET myelofibrosis, we further examined whether *CALR* mutation imparted similar proliferative and survival advantage to HSPCs of a patient who progressed to post-ET myelofibrosis (MF01, **Fig. 4g**). Interestingly, while mutant HSPCs showed increased cell cycle signature scores compared to wildtype HSPCs, mutant MkPs did not demonstrate increase in proliferative status compared to their wildtype counterparts (**Fig. 4h**), consistent with the patient’s clinical phenotype of a normal platelet count (318K/μL). Similar to our ET data, the earliest mutant HSPCs expressed higher anti-apoptotic signaling and *XBP1*-target genes, compared to the wildtype cells (**Fig. 4i**). Thus, these findings suggest enhanced survival of *CALR*-mutant HSPCs through upregulation of the IRE1-mediated UPR is maintained through disease progression to myelofibrosis.

### Multiplexed GoT reveals subclonal transcriptomic identities

Ongoing clonal evolution results in multi-clonal malignant populations following nested (linear) and non-nested (branched) structures. To define single-cell transcriptional program in relation to genotype thus often requires genotyping multiple mutational targets in parallel. To test the ability of GoT to target multiple genes in the same sample, we targeted three genes *CALR* (type 2 mutant, c. 1154_1155insTTGTC, VAF 43.5% by bulk exon sequencing), *NFE2* (c.782_785delAGAG, VAF 33%) and *SF3B1* (c.1984C>G, VAF 47.5%) in CD34^+^ cells sorted from peripheral blood mononuclear cells from a patient with myelofibrosis and anemia (MF02, **Fig. 5a**). Applying the pigeon-hole principle to the respective VAFs of these three mutations (as determined by targeted bulk sequencing in CD34^+^ cells), this malignancy has a clonal heterozygous mutation for *SF3B1,* a progeny subclone harboring a *CALR* mutation, which has an additional *NFE2* mutated progeny following a nested (linear) structure. GoT sequencing was performed on 8475 CD34^+^ cells, which provided targeted genotyping information for *CALR* and *NFE2* in 74% and 60% of cells, respectively.

**Figure 5.**
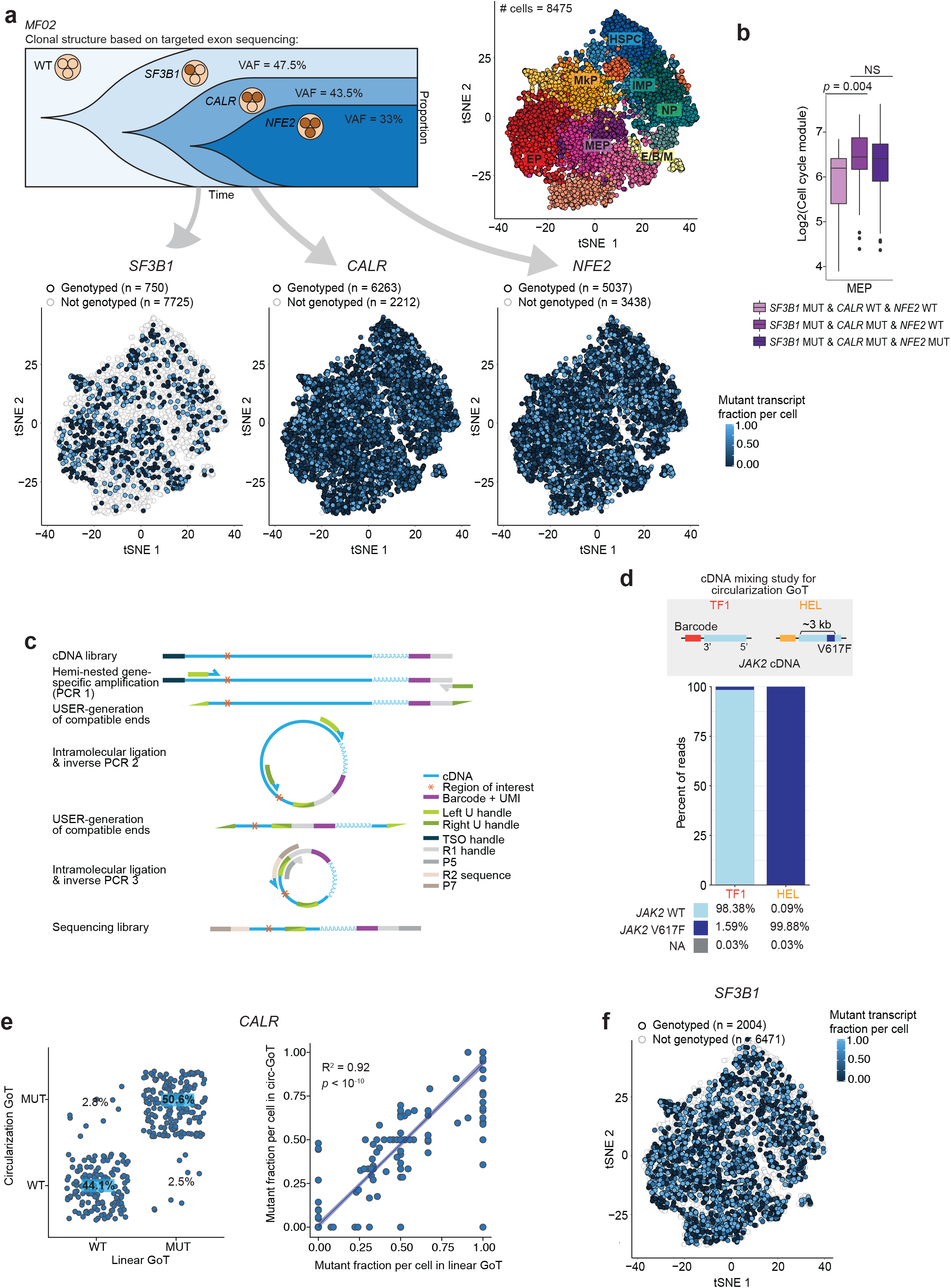
GoT can be extended to multiplexing to dissect subclonal identity and to targeting loci distant from the 3’ end via circularization. **(a)** Schematic of clonal evolution of neoplastic populations in patient MF02 based on targeted exon sequencing variant allele frequency (VAF). tSNE projection of CD34^+^ peripheral blood mononuclear cells using the Seurat package (see online methods). tSNE projections labeled with GoT information for *SF3B1*, *CALR* and *NFE2* mutations. **(b)** Cell cycle score in subclonal populations within the megakaryocytic-erythroid progenitor population (Wilcoxon rank-sum test). **(c)** Schematic representation of circularization GoT. **(d)** Mixing study with human *JAK2* wildtype (WT) cDNA from TF-1 cell line and homozygous *JAK2* V617F (MUT) cDNA from HEL cell line. Frequency of reads (WT, V617F or not assignable (NA)) that are assigned to TF-1 or HEL cell barcodes (CB). **(e)** Comparison of genotype assignment for *CALR* in patient sample MF01 between linear GoT and circularization GoT after downsampling reads to 300K with 10 iterations (left). Comparison of *CALR*-mutant UMI fraction per cell between GoT and circularization GoT (circ-GoT) (two-tailed Pearson’s correlation, right). **(f)** tSNE projection of CD34^+^ cells from sample MF02 labeled with GoT information for *SF3B1* from circularization and linear GoT. HSPC, hematopoietic stem progenitor cells; IMP, immature myeloid progenitors; NP, neutrophil progenitors; E/B/M, eosinophil, basophil, mast cell progenitors; MEP, megakaryocytic-erythroid progenitors; MkP, megakaryocytic progenitors; EP, erythroid progenitors; WT, wildtype; MUT, mutant.

In this context, GoT allows to compare the transcriptional outputs of the different mutation alone or in combination. For example, since *SF3B1* mutations have been shown to block erythroid maturation^48^, we examined whether the addition of a *CALR* mutation would still confer increased proliferative status in MEPs. We found that *SF3B1*/*CALR*-double mutants exhibited increased proliferative advantage over *SF3B1*-single mutants (**Fig. 5b**). In contrast, we found that the addition of *NFE2* mutation (triple-mutant) did not increase cell cycle activation beyond the activation seen in the *SF3B1/CALR* double mutant, unlike *in vitro* data that suggested that *NFE2* mutation in conjunction with *JAK2* V617F may enhance the proliferative advantage of mutant cells^49^. In summary, multiplexed GoT demonstrated the ability to interrogate complex clonal structures, as well as the need to assess the combinatorial transcriptional output of mutations in the context of high-resolution cell identity mapping.

### Circularization GoT enables genotyping of mutations distant from the 3’ end

GoT amplicon recovery of a mutation that is distant from the 3’ end (>1.5 kb, e.g. *SF3B1* genotyping of 9% of cells, **Fig. 5a**), was lower than for targets closer to the 3’ end such as *CALR* variants (~800 bp) and *XBP1* splice site (1.3 kb). While analysis of driver mutations revealed that the majority of targets are within 1.5 kb of either the 3’ or 5’ ends, and thus amenable to GoT with either the 3’ or 5’ procedures (**Supplementary Fig. 9**), other loci of interest reside 1.5- 3 kb from either end, and thus the dependency on relative proximity to the 3’ or 5’ end is limiting. We reasoned that the lower efficiency resulted, at least partially, from the inability of larger targeted amplicon fragments to cluster efficiently on the flow cell during sequencing. To overcome this limitation and to allow universal GoT for all expressed genes, we developed a method in which sequential rounds of circularization and inverse PCR remove the intervening sequence between the region of interest and the barcodes (**Fig. 5c**). This method generates amplicons that retain contiguity of the original molecules but are short enough to cluster effectively to be sequenced with standard parameters.

In order to demonstrate the ability of circularization GoT to maintain the CB with the correct genotyping segment despite significant distance from the 3’ end, we targeted *JAK2* V617F, which is ~3 kb away from the 3’ end of the transcript. We performed circularization GoT on mixed cDNA sample in which barcoded cDNA from TF1 cell line with wildtype *JAK2* was mixed with barcoded cDNA from HEL cells with homozygous *JAK2* V617F (**Fig. 5d, Supplementary Fig. 10a**). Reads with the expected primer sequences provided highly specific genotyping assignment, such that 98.38% of reads with the TF-1 CB was associated with the correct wildtype *JAK2* sequence and 99.88% of reads with the HEL CB with the mutant *JAK2* sequence. Thus, this experiment demonstrated the ability of circularization GoT to target mutations, including SNVs, distant from the 3’ end. We further validated this method by comparing the genotyping information to the previously described un-circularized (linear) GoT data for *CALR* genotyping, and found strong concordance between the genotyping results of the two methods (**Fig. 5e, Supplementary Fig. 10b,c**). Finally, we applied circularization GoT to the *SF3B1* mutation target from sample MF02, which significantly increased the yield of genotyped cells from 750 cells to 2004 cells (9% to 24% of total cells, **Fig. 5f**). Thus, these results demonstrate the ability of circularization GoT to provide single cell genotyping in patient samples in a highly specific and efficient manner, extending its reach even to targets distant from gene ends.

## DISCUSSION

High-throughput digital scRNA-seq methods have been applied comprehensively to the study of human cancer samples to reveal significant intra-tumoral diversity, due to their ability to profile many thousands of cells at lower cost per cell. Nevertheless, these studies have been limited by the inability to couple genotypic information (other than large copy number aberrations) with single-cell whole transcriptome profiles, thereby precluding the ability to distinguish accurately between malignant and wildtype cells or between different subclones. While full transcriptome single-cell sequencing was shown to enable capture of genotypes at lower efficiency^50^ or with targeted amplification (e.g. *BCR-ABL* rearrangement)^15^, these methods are limited in throughput. To overcome this limitation, we present here GoT which capitalizes on high-throughput scRNA-seq (in this case, the 10x Genomics Chromium single cell 3’ platform), by which thousands of cells can be jointly profiled for genotyping information as well as single-cell full transcriptomes.

MPNs serve to highlight the need for high-throughput joint single-cell genotype and whole transcriptome technology. Mutated cells are not readily distinguishable from the wildtype progenitor cells based on surface markers. Furthermore, recent studies of normal hematopoiesis demonstrated the significant complexity of the differentiation topology, which requires the throughput of thousands of single-cells for accurate mapping^51-53^. Indeed, since the discovery of *CALR* mutations in MPN patients in 2013, many studies have attempted to characterize the underlying mechanisms by which mutations in a chaperone protein like CALR result in MPNs^11,26^. While the importance of its association with MPL has been well-established by several groups^25,29,30,54^, other studies have yielded conflicting findings regarding the downstream pathways in *CALR*-mutated cells^28,55,56^. These conflicting reports may result in part from the inherent limitations of alternative methods, which use *in vitro* culture or mouse modeling.

GoT allowed us to directly interrogate the transcriptional impact of *CALR* mutation in HSPCs and progenitor cells from primary bone marrow samples from ET patients. In particular, wildtype cells found in the same samples provide an ideal comparison data set, controlling for patient-specific and technical confounders. Through the application of GoT, we further isolate the impact of *CALR* mutations on each progenitor subset. We observed that mutant *CALR* provided a greater fitness advantage through differentiation, associated with higher cell-cycle gene expression in committed progenitors (most pronounced in MEPs and MkPs) compared with uncommitted HSPCs. In addition, the ability to fine-map mutant vs. wildtype transcriptional differences in HSPCs revealed upregulation of NF-κB pathway genes in the most undifferentiated HSPCs. We further applied GoT to target the unconventional splice site of *XBP1* to demonstrate that IRE1 activity was increased in the HSPC mutants. Our data thus nominates IRE1/XBP1 pathway as a potential therapeutic target for eradication of the mutant clone in the uncommitted HSPCs in patients with *CALR*-mutated MPNs.

Clonal evolution with acquisition of additional somatic mutations is one of the hallmarks of cancer including myeloid neoplasms. Therefore, the ability of GoT to genotype multiple target genes in parallel is critical. Moreover, while described here for 3’ droplet-based scRNA-seq, GoT can be integrated in any scRNA-seq method that generates full length cDNA as an intermediate product (e.g. Microwell-seq^57^, 10x Single Cell V(D)J + 5′ Gene Expression). GoT can also be readily integrated with methods that layer additional measurements on top of transcriptomes (e.g. protein levels via CITE-seq^58^) and methods for multiplexing samples in high-throughput scRNA-seq (e.g. cell hashing^59^ and related methods^60,61^). While this method is limited to genotyping of transcribed mutations, our analysis shows that most driver mutant alleles are transcribed and may even be expressed preferentially. Thus, GoT delivered precision genotyping even in the highly challenging setting of human CD34^+^ progenitor cells, where the magnitude of transcript per cells is significantly lower than in epithelial cancers^62^.

In conclusion, high-throughput linking of single-cell genotyping of expressed genes to transcriptomic data may provide the means to gain insight into questions such as the integration of clonal diversification with lineage plasticity^63^ or differentiation topologies^3,50^. This method will also likely be a valuable tool to assess the impact of somatic mutations in pre-malignant states (e.g., clonal hematopoiesis of indeterminant potential), in which mutant and wildtype cells are phenotypically similar, requiring higher resolution single-cell transcriptional interrogation. Thus, GoT may pave the way to resolve central questions relating to the link between genetic mutations and cellular identities, and to unravel the underlying programs that enable clonal expansions and evolution in human neoplasms.

## ONLINE METHODS

### Species mixing experiment

Previously published UT7 and Ba/F3 cell lines expressing human *MPL* and either human wildtype *CALR* or mutant *CALR* (type 1, 52 bp deletion) were used for the species mixing study^25^. Briefly, human *MPL* expressing Ba/F3 and UT-7 cell lines were generated by retroviral transduction, after which they were subjected to infection with *CALR* variant lentiviral supernatants. Wildtype UT7 cells and mutant Ba/F3 cells were mixed in equal proportions which underwent GoT, targeting ~1000 cells.

### Patient samples

Cryopreserved bone marrow aspirates (BMAs) or peripheral blood mononuclear cells (PBMCs) from patients with documented *CALR* mutations which had been banked under specimen acquisition and molecular profiling protocols approved by the Institutional Review Board of Memorial Sloan-Kettering Cancer Center were retrieved after a database search. See **Supplementary Table 1** for clinical information. All patients provided informed consent. Cryopreserved BMAs or PBMCs were thawed and stained using standard procedures (10 min, 4°C) with the surface antibody CD34-PE-Vio770 (clone AC136, Miltenyi Biotec) and DAPI (Sigma-Aldrich). Cells were then sorted for DAPI-negative, CD34^+^ and CD34-negative cells using BD Influx at the Weill Cornell Medicine flow cytometry core.

### Targeted myeloid panel

To determine the presence and location of recurrent somatic mutations and their VAF, targeted next-generation sequencing was performed on DNA samples extracted from frozen unfractionated PBMCs (patients MF01), CD34-negative sorted BMAs (patients ET02 and ET03), CD34^+^ sorted BMA (patient ET01) and CD34^+^ sorted PBMC (patient MF02), as previously described^64^. Briefly, targeted enrichment of 45 genes (*ABL1, ASXL1, BCOR, BRAF, CALR, CBL, CEBPA, DNMT3A, ETV6, EZH2, FAM5C, FLT3, GATA1, GATA2, HNRNPK, IDH1, IDH2, IKZF1, JAK1, JAK2, KDM6A, KIT, KRAS, MPL, NFE2, NOTCH1, NPM1, NRAS, PHF6, PTPN11, RAD21, RUNX1, SETBP1, SF3B1, SH2B3, SMC1A, SMC3, SRSF2, STAG2, SUZ12, TET2, TP53, U2AF1, ZRSR2*) recurrently mutated in myeloid malignancies was performed using the Thunderstorm system (Raindance Technologies, Billerica, MA) using a custom primer panel followed by sequencing using the Illumina MiSeq (v3 chemistry).

### Droplet digital PCR

Peripheral blood from three ET patients with mutations in *CALR* underwent Ficoll density gradient separation, immunomagnetic selection for CD34+ cells (Miltenyi Biotech), and fluorescence-activated cell sorting (FACS) (Influx, Becton-Dickinson) using PeCy7-labeled CD34, clone 561 (Biolegend) and APC-labeled CD38, clone HIT2 (Biolegend) antibodies to isolate CD34^+^CD38^-^ and CD34^+^CD38^+^ cell compartments. DNA was extracted from sorted cells (Qiagen) and the variant allele frequency (VAF) of *CALR* mutations was measured by droplet digital PCR (QX200 Droplet Digital PCR System, Bio-Rad) with primers that specifically detect *CALR* type 1 mutations (52-bp deletion (p.L367fs*46), *CALR* type 2 mutations (5-bp TTGTC insertion (p.K385fs*47) or normal alleles.

### Genotyping of Transcriptomes (GoT)

The standard 10x Genomics Chromium v.2 protocol was carried out according to manufacturer’s recommendations (10x Genomics, Pleasanton, CA) until after emulsion breakage and recovery of first strand cDNA (**Fig. 1a,** step 1). If the targeted gene of interest, e.g. *SF3B1*, was not robustly detected by the standard 10x procedure (i.e. if <60% of the expected cells showed expression) based on *a priori* knowledge in a similar dataset, a gene-specific primer was spiked into 10x primer mix at 1% of the concentration of the cDNA amplification primers for the initial cDNA PCR step (**Fig. 1a,** see **Supplementary Table 3** for list of primers). Then, during the amplification step, the 10x cDNA library underwent an extra cycle of PCR beyond the manufacturer’s recommended number of cycles. After cleanup with SPRIselect, a small portion of the cDNA library, 3 μL (~10% of total) was aliquoted for targeted genotyping, and the remaining cDNA underwent the standard 10x protocol. The cDNA set aside for GoT was amplified for 3 to 4 additional cycles using KAPA HiFi HotStart ReadyMix (KAPABiosystems) and 10x primer mix to provide sufficient material for the enrichment step. After clean-up, locus-specific reverse primers and the generic forward SI-PCR were used to amplify the site of interest of the cDNA template (**Supplementary Table 3**) using ~10 PCR cycles (**Fig. 1a**). The locus-specific reverse primers contain a partial Illumina read 2 handle, a stagger to increase the complexity of the library for optimal sequencing and a gene specific region to allow specific priming. The SI-PCR oligo (10x) anneals to the partial Illumina read 1 sequence at the 3’ end of the molecule, preserving the cell barcode (CB) and UMI. After the initial amplification and SPRI purification to remove unincorporated primers, a second PCR was performed with a generic forward PCR primer (P5_generic) to retain the CB and UMI together with an RPI-x primer (Illumina) to complete the P7 end of the library and add a sample index. The targeted amplicon library was subsequently spiked into the remainder of the 10x library to be sequenced together on a HiSeq 2500 or sequenced separately on MiSeq with v3 chemistry (Illumina). The cycle settings were as follows: 26 cycles for read 1, 98 or 130 cycles for read 2, and 8 cycles for sample index.

### Single-cell RNA-seq data processing, alignment, cell type classification and clustering

10x data was processed using Cell Ranger 2.1.0 with default parameters. Reads were aligned to the human reference sequence GRCh38 or hg19 or to mouse reference mm10 (species mixing experiment). The genomic region of interest for genotyping was examined to determine how many UMIs with targeted sequence were present in the conventional 10x data (**Fig. 1c, Supplementary Fig. 4**). The Seurat package was used to perform unbiased clustering of the CD34^+^ sorted cells from patient samples^65^, following the publicly available Guided Clustering Tutorial. Briefly, cells with UMI <200 or UMI > 3 standard deviation from the mean UMI and mitochondrial gene percentage >10% were filtered. The data was log normalized using a scale factor of 10,000. Potential confounders (e.g., number of UMI per cell and the proportion of mitochondrial genes) were regressed out of the data before principle component analysis (PCA) was performed using variable genes. JackStraw method was used to determine the statistically significant PCs to be used for graph-based clustering. t-SNE was used to visualize the clusters. Clusters were manually assigned based on differentially expressed genes using the FindAllMarkers function. Definitions of CD34^+^ progenitor subgroups from Velten *et al* (2017) were used as a reference to assign the clusters (**Supplementary Fig. 7a**)^17^. Pseudotime analysis was performed using the Monocle R package^33^.

### IronThrone GoT: targeted genotype amplicon sequence processing and mutation calling

To ensure correct priming, targeted amplicon reads (read 2) were screened for the presence of the primer sequence and the expected intervening sequence between the primer and the start of the mutation site (‘shared sequence’, **Supplementary Fig. 2a,b**). 90.0% of the reads from the mixing study showed the expected primer and shared sequences. Subsequently, for reads that passed the priming step, the corresponding read 1 was screened for the presence of the 16 bp long CB that matched the CB in the whitelist provided by 10x Genomics (**Supplementary Fig. 11**). For CB reads that were 1-Hamming-distance away from the whitelisted CB, the probability that the observed barcode originated from the whitelisted CB was calculated taking into account the base quality (BQ) score at the differing base. The whitelisted CB with the highest probability was used to replace the observed CB, only if the probability exceeded 0.99. For the duplicate reads with the same CB and UMI, the genotype (wildtype vs. mutant) of the read with the highest BQ score was assigned for the given UMI.

The species-mixing study for *CALR* mutation was used to further optimize the pipeline for mutation calling. We integrated targeted amplicon measures including BQ, number of base pair mismatches, and number of duplicate reads per UMI, and determined optimized parameters that maximize the number of genotyped cells while minimizing genotype mis-assignment (**Supplementary Fig. 3a-e**). Setting thresholds for the minimum number of duplicate reads and maximum frequency of mismatches contributed significantly to filtering out mis-assigned reads likely due to PCR recombination/errors. A combination of threshold of two or more duplicate reads for a given UMI and a threshold of allowing less than or equal to 0.2 mismatch ratio significantly improved correct assignment of cells, while maximizing the number of included cells for analysis, and was adopted in the analysis here (**Supplementary Fig. 3a**). Results of the precision and recall analyses also affirmed this combination of thresholds for minimum duplicate reads and maximum mismatch ratio (**Supplementary Fig. 3b**). Moreover, given the high number of *CALR* transcripts in the cell lines and thus high PCR recombination rate, cells were assigned as wildtype or mutant if >90% of *CALR* amplicon UMIs were wildtype or mutant, respectively.

To further assess the impact of various parameters of the amplicon reads on the precision of mutation calling, we tested these parameters in a random forest classification using the mixing study, as implemented in the R randomForest package^66^ (**Supplementary Fig. 3c-e**). Mean decrease accuracy was determined as a measure of importance of each variable used for the calculation of splits in trees (**Supplementary Fig. 3c**). For each combination of mismatch ratio and duplicate thresholds, random forest was run 100 times to find the optimal number of random variables used in each tree and the minimum out-of-bag error was selected (**Supplementary Fig. 3d,e)**. This random forest analysis also showed a minimum duplicate read threshold of 2 and maximum mismatch ratio threshold of 0.2 to be optimal for minimizing mis-assignments, and the relatively low contribution of additional quality metrics (**Supplementary Fig. 3c-e**).

The genotyping information is derived from transcribed molecules and may be affected by the capture of transcripts from wildtype vs. mutant alleles of heterozygous mutations in primary patient samples. This may be due to incomplete sampling of the transcript pool or due to transcriptional bursts, which leads to skewed transcript pools. Consequently, as the number of UMIs per cell increases, the likelihood of capturing a mutant transcript increases, resulting in an apparently higher frequency of mutated cells. Thus, the number of mutant reads may be underestimated in cells with lower amplicon UMI counts. Nonetheless, the frequency of mutant cells (e.g., 26% in patient sample ET01) as determined by GoT using all cells that harbor at least one UMI yielded values that were similar to that determined by bulk DNA exon sequencing of *CALR* from CD34^+^ cells (mutant cell fraction of 30% based on VAF of 0.15 in a diploid heterozygous mutation), suggesting that in this context the mutant allele may be preferentially expressed, limiting biases resulting from small UMI numbers and incomplete sampling of heterozygous mutations.

While the bulk of the downstream analyses between *CALR* mutant and wildtype cells used a threshold of two or greater genotyping amplicon UMIs, we systematically applied three approaches to exclude the impact of this confounder (i.e., expression level of target gene) on the conclusions. First, to exclude the possibility that higher *CALR* expression in committed progenitors can result in greater ability to detect mutant alleles and thereby result in higher mutated cell frequency, we down-sampled all cells to a single amplicon UMI prior to mutation calling and found that the increase in mutation frequencies in MkP compared with HSPC remained unchanged (**Fig. 2d**). Second, we explored the sensitivity of the difference between mutant and wildtype cells (e.g. cell cycle module score) by increasing the minimal amplicon UMI threshold allowed for mutation calling and demonstrated that this did not impact the central findings of this study (**Supplementary Fig. 5a-c**). Third, we explicitly modeled the impact of *CALR* amplicon UMI in multivariable models (generalized linear model using R Stats package v3.5.1, e.g., pseudotime analysis), in which the number of amplicon UMI was included in the model alongside the mutation status (**Fig. 2f**).

### Differential expression and gene set enrichment analysis

For gene module analysis, the aggregate gene expression levels of modules of genes involved in biological processes of interest (see complete list of genes for each module in **Supplementary Table 2**) were measured by calculating the ratio of the sum of the UMI of genes within the module over total UMI per cell, scaled to 10,000. For boxplots, log2(expression +1) is shown. All of the gene modules have been previously published (see references in main text), except for the MkP and HSPC1 module, which are composed of genes that were highly expressed in these respective populations. Differential gene expression analysis between mutant and wildtype cells for each of the progenitor cluster for each patient was performed via the ‘FindMarkers’ function within the Seurat Package using the Tobit-test for differential gene expression^33^. The differentially expressed genes were examined individually for each patient; they were also examined in combination for each cluster across the patients by combining the *P*-values for the differentially expressed genes via Fisher’s method and averaging the log2(fold change). Genes that were differentially expressed with FDR <0.25 were included for gene set enrichment analysis. Hypergeometric test for gene set enrichment analysis was performed using the gProfiler package^67^. KEGG and Reactome data sources were included in the analyses.

### Circularization GoT

For patient samples, we used the same starting material as for GoT (i.e. re-amplified cDNA fraction from 10x library); for the *JAK2* cDNA mixing study, we mixed barcoded cDNA from two cells lines (TF1: *JAK2* WT, HEL: Homozygote *JAK2* V617F). With these cDNA libraries, we first performed a PCR to enrich for an amplicon from ~50 bp upstream our region of interest to the 3’ end of the 10x library fragment, therefore retaining cell barcode (CB) and unique molecule identifier (UMI), using KAPA HiFi Uracil+ master mix (Kapa Biosystems) and the following PCR conditions: 98°C for 45”; 10 to 20 cycles of: 98°C for 20”, 65°C for 30”, 72°C for 2’, 72°C for 5’. Complementary U-overhang are added to the forward (Fw) and reverse (Rv) primers to allow circularization: Fw-primer#1: AGGUCAGTCU-[50bp-upstream-locus-specific], Rv-primer#1: AGACUGACCUCTACACGACGCTCTTCCGATCT (**Supplementary Table 3**). For gene lowly represented in the cDNA library (such as *SF3B1*), we specifically pre-enriched the locus of interest by doing a PCR targeting ~100bp upstream our region of interest to the 3’ end of the 10x library fragment, using KAPA HiFi Ready mix (Kapa Biosystems) and the following PCR conditions: 98°C for 45”; 20 cycles of: 98°C for 20”, 67°C for 30”, 72°C for 2’, 72°C for 5’. PCR product resulting from the first single or double PCR was then cleaned-up and concentrated using 1.3X SPRI beads. Next, amplicon cohesive ends were created using 40U/mL USERII enzyme (M5508-NEB) digestion for 1 hour at 37°C in 1X CutSmart buffer. Reaction was stopped by incubating for 10’ at 65°C. Relying on complementary overhang at both end of the amplicon, circularization was performed in a large volume (>1 mL) to favor intra-molecule ligation. The following reaction was set-up and incubated overnight at 16°C: USERII-digested amplicon, 2000 U/mL T4 ligase (NEB), 1X CutSmart Buffer (NEB), 1 mM ATP (Roche). Next, T4 DNA ligase was inactivated by incubating for 15’ at 70°C. Then, unwanted un-ligated products were removed by adding 6U of lambda exonuclease (NEB, M0262S) in the ligation mix and incubating for 30’ at 37°C. Exonuclease was inactivated for 20’ at 65°C. Ligated product was cleaned-up and concentrated using 1.3X SPRI beads. A second PCR was set-up to retain the locus of interest and barcodes on the same molecule while removing the unwanted 3’ downstream region of the targeted region. PCR reaction was set-up and performed as previously described using the following primers: Fw-primer#2: AGGUCAGTCU[3’end-locus-specific], Rv-primer#2: AGACUGACCU[10bp-downstream-locus-specific]. After PCR#2, SPRI clean-up, USERII digestion, overnight T4 ligation, lambda exonuclease digestion were performed as previously described. After the second ligation, the ligated product was again cleaned-up and concentrated using 1.3X SPRI beads. To linearize the product of ligation, we performed a third PCR: Fw-primer#3: CCTTGGCACCCGAGAATTCCA[10bp-upstream-specific-locus], Rv-primer#3: SI-PCR (10x Genomics). We used KAPA HiFi master mix (Kapa Biosystems) and the following PCR conditions: 98°C for 45”; 10 cycles of: 98°C for 20”, 58°C for 30”, 72°C for 1’; 72°C for 5’. After SPRI purification, a last PCR was performed with a generic forward PCR primer (P5_generic) and an RPI-x primer (Illumina) to complete the P7 end of the library and add a sample index. The targeted amplicon library was subsequently sequenced using PE150 on MiSeq (Illumina).

### Comparison of mutant allelic fraction in whole exome sequencing (WES) and RNA-seq

We compared the mutant allelic fractions between gDNA and RNA, estimated from WES and RNA-seq data, respectively, in five cancer cohorts (BRCA, breast invasive carcinoma; HNSC, head and neck squamous cell carcinoma; KIRC, kidney renal clear cell carcinoma; LUAD, lung adenocarcinoma; STAD, stomach adenocarcinoma). For this analysis, we thank Dr. Tae-Min Kim (Cancer Research Institute, College of Medicine, The Catholic University of Korea) for sharing the curated datasets based on his previous study^68^. In brief, the datasets of each cancer cohorts were initially prepared with somatic mutation sets that are reported from The Cancer Genome Atlas (TCGA) portal (https://portal.gdc.cancer.gov/). Then, reference and alternative alleles for these mutations were counted in bam files of WES and RNA-seq using SAMtools mpileup^69^, and filtered for >10 coverage of reads. We then converted genomic coordinates of the datasets from hg19 to hg38 assembly. To identify the frequencies of somatic mutations in cancers, we used the CNT values from CosmicCodingMuts.vcf (v86) in the Catalogue of Somatic Mutations in Cancer (COSMIC) database^70^. Then, we further annotated the variants as oncogene, tumor suppressor gene, (TSG) or passenger (Vogelstein *et al*. 2013^71^ and Bailey *et al*. 2018^72^).

### Determination of targeted loci distance from 3’ or 5’ ends of transcripts

To identify Ensembl transcript ID (ENST) corresponding to each mutation in the datasets of five cancer cohorts described above, we matched them with COSMIC ID and annotated from the file of CosmicMutantExport.tsv (v86). We used the biomaRt R package^73^ with the GRCh38 version to annotate the transcript including the length of transcript and the position of cDNA start codon in the transcript. The positions of the 5’ untranslated region (UTR) ends were determined to calculate the distance from 5’ end to target site.

### Data availability

All of the sequencing data and the IronThrone GoT pipeline will be available upon publication.

## ACKNOWLEDGMENTS

We thank Dr. Ann Mullally (Brigham and Women’s Hospital) for sharing the cell lines for the species-mixing study. This work was funded by the Columbia University Physical Sciences in Oncology Center Pilot Grant. R.C. is supported by Lymphoma Research Foundation and Marie Skłodowska-Curie fellowships. D.A.L. is supported by the Burroughs Wellcome Fund Career Award for Medical Scientists, the ASH Scholar Award, Leukemia Lymphoma Society Translational Research Program, the National Institutes of Health (NIH) Big Data to Knowledge initiative (BD2K, 1K01ES02543101), and the Stand Up To Cancer Innovative Research Grant (SU2C-AACR-IRG-0616). Stand Up To Cancer is a program of the Entertainment Industry Foundation. Research grants are administered by the American Association for Cancer Research, the scientific partner of SU2C. The work was enabled by the Weill Cornell Epigenomics Core and Flow Cytometry Core.

---

**Supplementary Figure 1.**
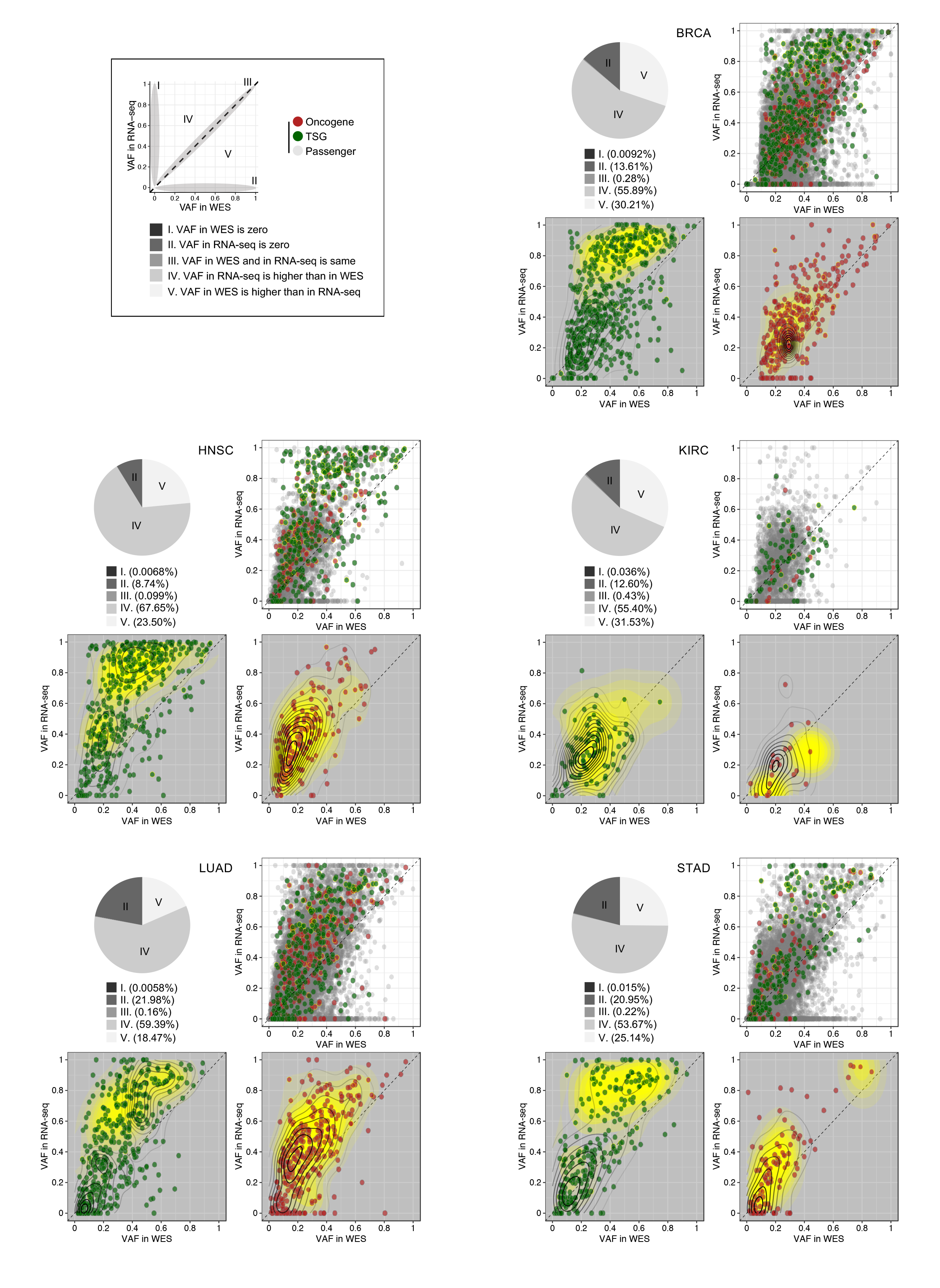
Comparison of variant allele frequency (VAF) between whole exome sequencing (WES) and RNA-seq.

Pie charts show the fraction of variants that are categorized as described in the top-left-box. Distribution of mutant allele fraction is annotated as oncogene, tumor suppressor gene (TSG), and passenger mutations in the right panel (definitions according to Vogelstein *et al*. 2013^71^ and Bailey *et al*. 2018^72^). Diagonal dashed lines indicate equal allelic fraction between WES and RNA-seq. Yellow density contours represent driver distributions. BRCA, breast invasive carcinoma; HNSC, head and neck squamous cell carcinoma; KIRC, kidney renal clear cell carcinoma; LUAD, lung adenocarcinoma; STAD, stomach adenocarcinoma.

**Supplementary Figure 2.**
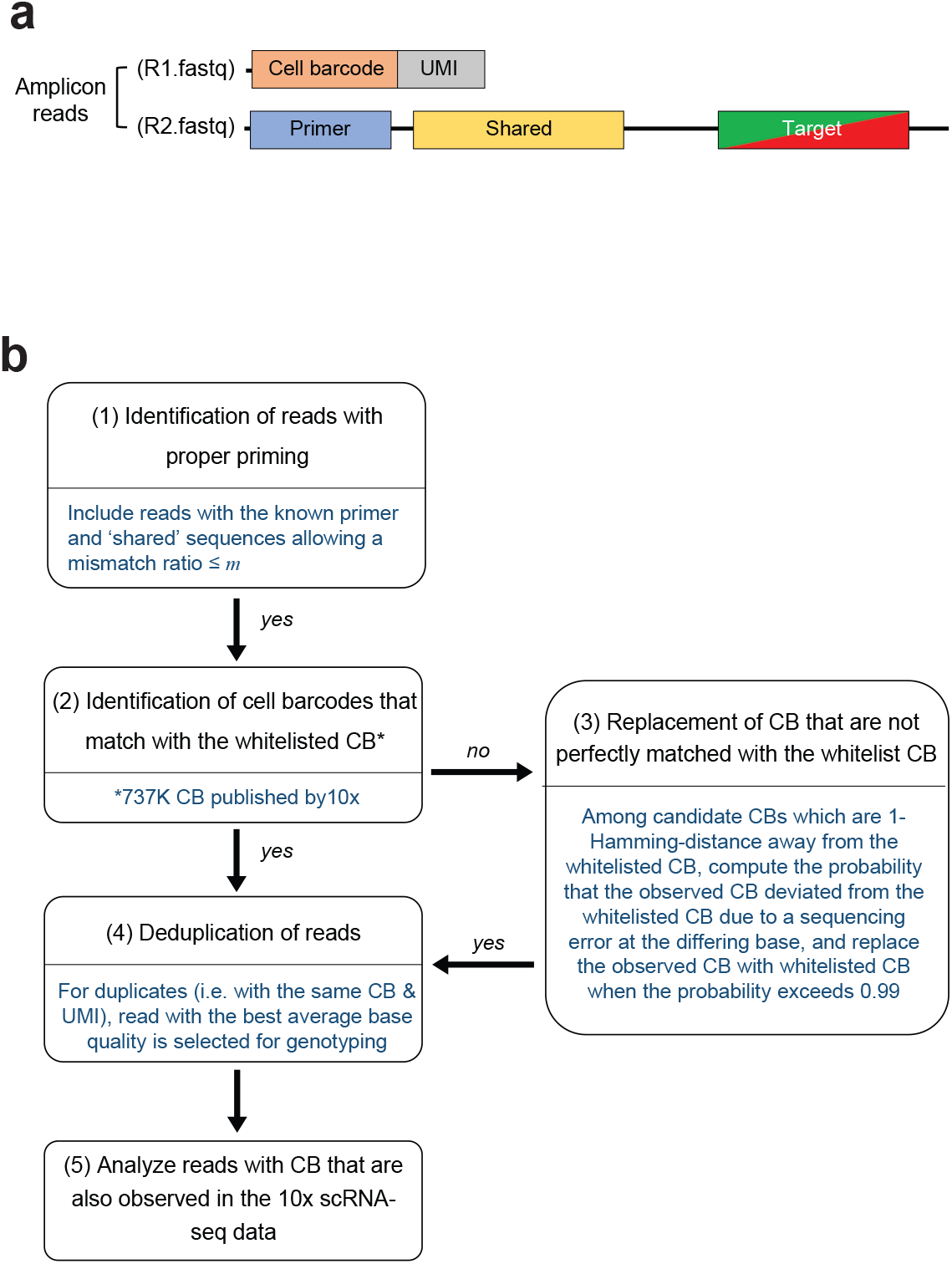
Analysis pipeline of Genotyping of Transcriptomes (GoT). **(a)** Simplified representation of amplicon reads. **(b)** Flow chart of the GoT analysis pipeline (see online methods).

**Supplementary Figure 3.**
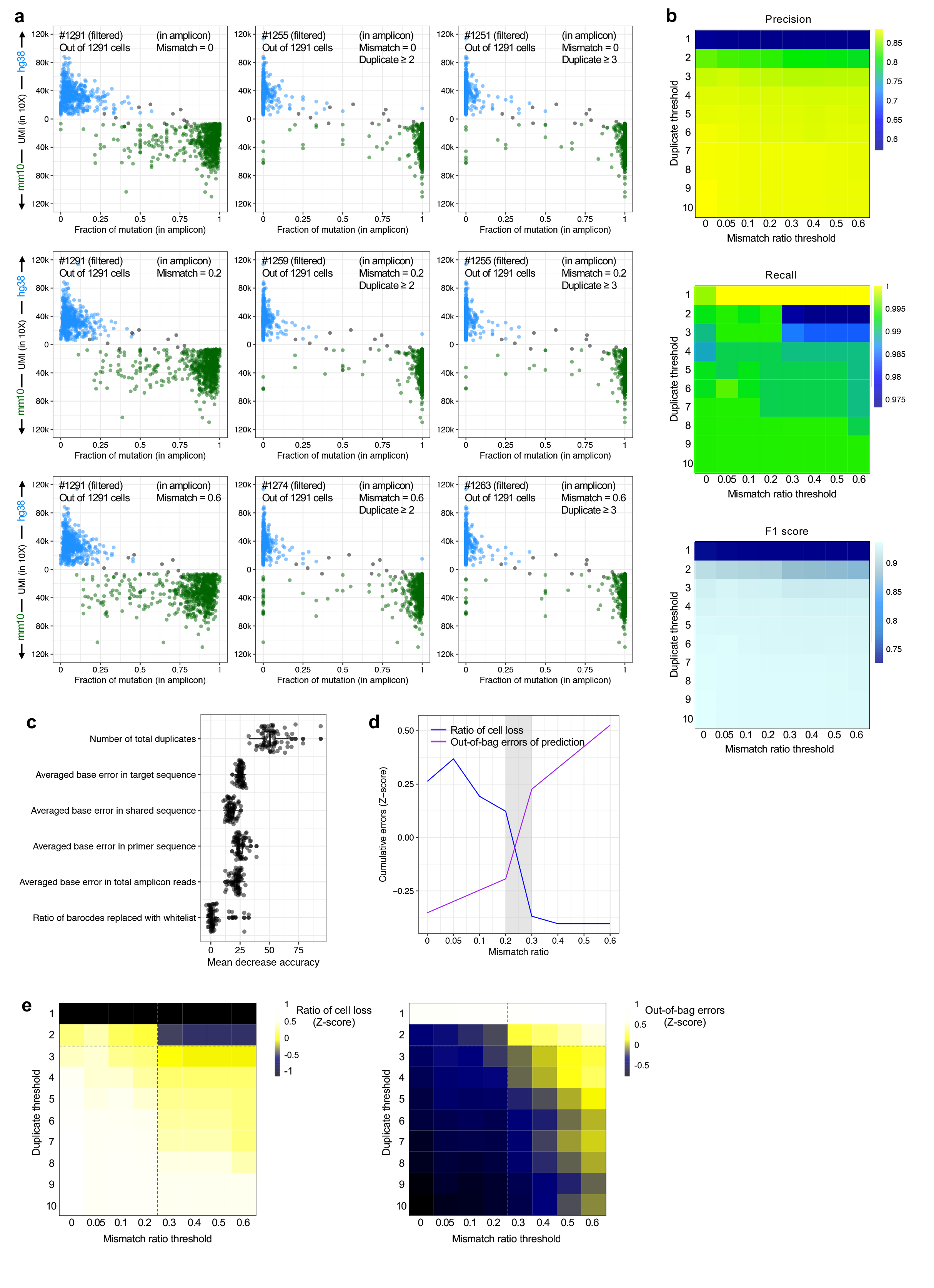
Optimization of parameters in targeted amplicon sequence processing pipeline in Genotyping of Transcriptomes (GoT). **(a)** Murine (green) vs. human (blue) genome alignment of 10x data (y-axis) with genotyping data by GoT (x-axis) with various thresholds for minimum duplicate reads (across) and maximum mismatch ratio (down). Multiplet cells are labeled in gray. **(b)** Results of precision, recall and F1 score analysis for combinations of minimum duplicate reads and maximum mismatch ratios. **(c)** Measure of importance of each variable used for the calculation of splits in trees in random forest classification test. **(d)** Ratio of cell loss and genotyping errors (Z-score in y-axis) based on mismatch ratio thresholds (x-axis); area of intersection is highlighted with gray around the mismatch ratio 0.2. **(e)** Heatmaps showing Z-scores of number of filtered cells (left) and predicted error rates (right) from random forest classification tests for combinations of minimum duplicate reads and maximum mismatch ratio thresholds.

**Supplementary Figure 4.**
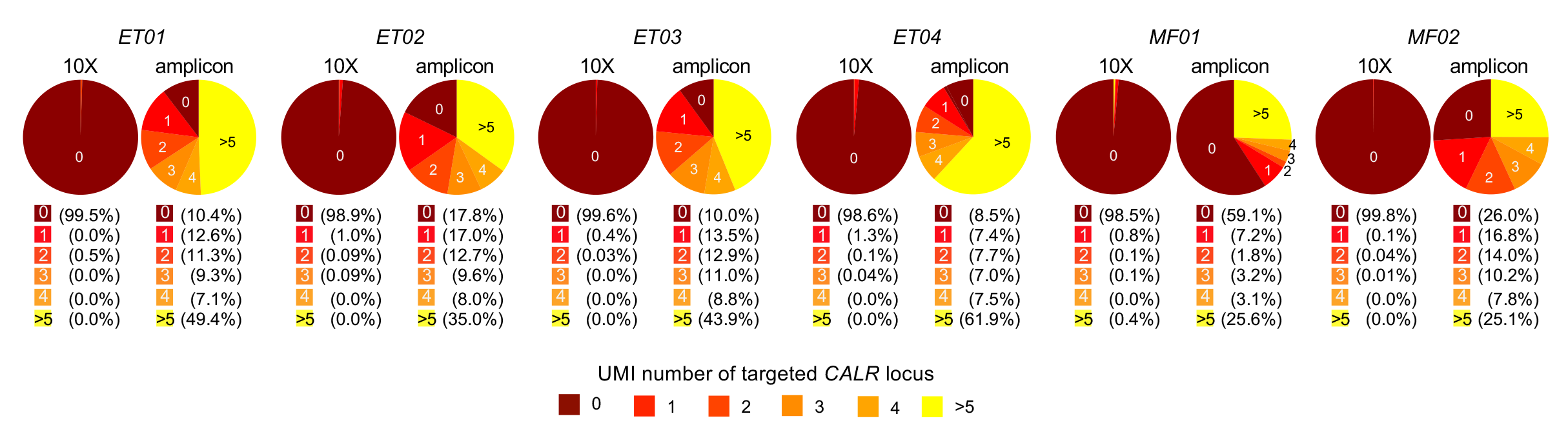
GoT captures genotyping information of single cells through cDNA. Percentage of cells with *x* number of UMIs with *CALR* mutation locus in 10x data (left) and GoT data (right).

**Supplementary Figure 5.**
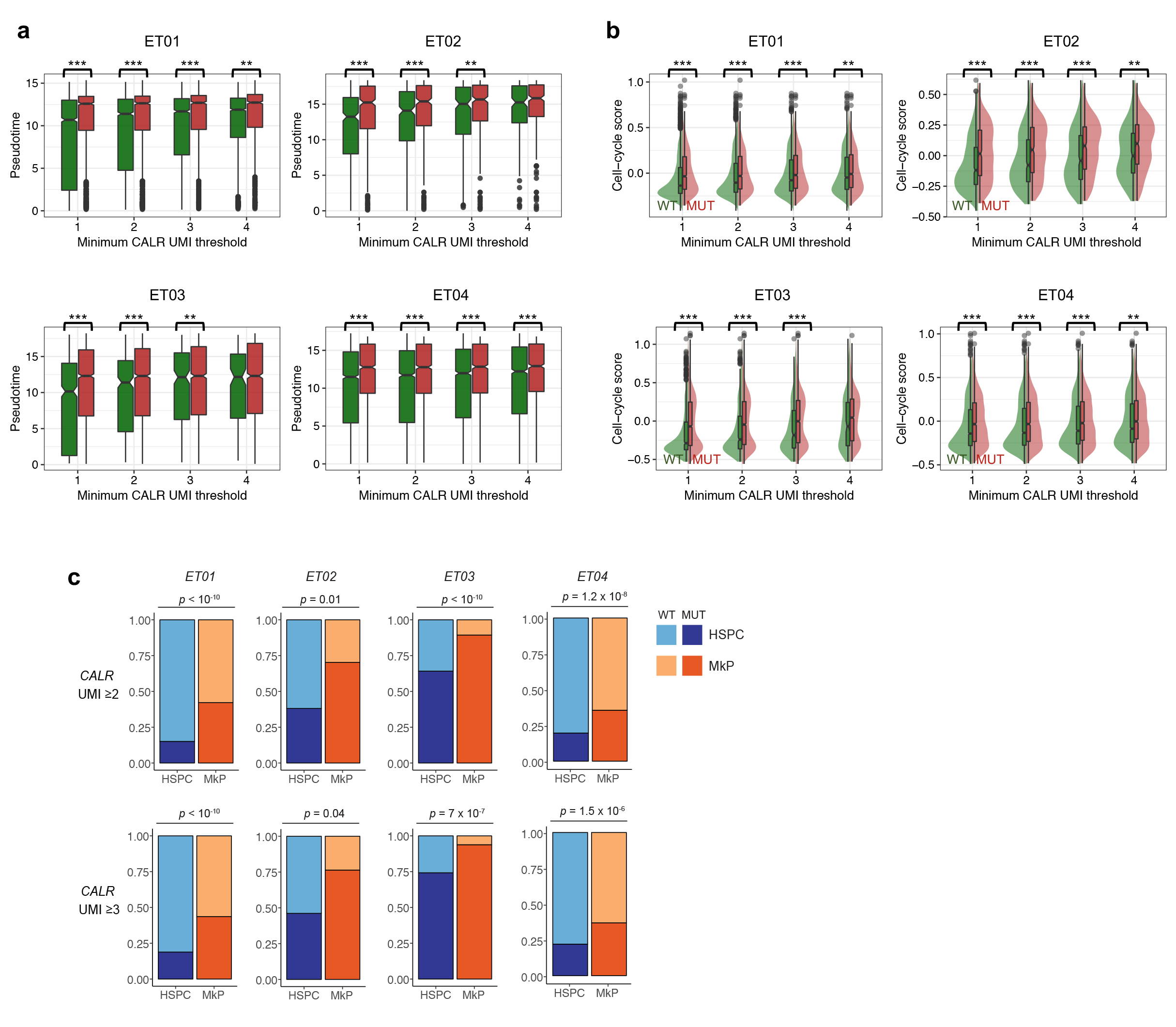
Results of GoT analysis is robust to various amplicon UMI thresholds. **(a)** Pseudotime difference between wildtype (WT) and mutant (MUT) cells with increasing number of thresholds for targeted genotyping UMI. \*\**P* < 0.01; \*\*\**P* < 0.001 (two-tailed t-test). (**b**) Cell cycle module score between WT and MUT cells with increasing number of thresholds for targeted genotyping UMI. \*\**P* < 0.01; \*\*\**P* < 0.001 (two-tailed t-test). **(c)** WT and MUT cell frequency in hematopoietic stem progenitor cell (HSPC) (blue) vs. megakaryocytic progenitor (MkP) (orange) clusters with a minimum genotyping UMI threshold of 2 (top) and 3 (bottom, Fisher’s exact test).

**Supplementary Figure 6.**
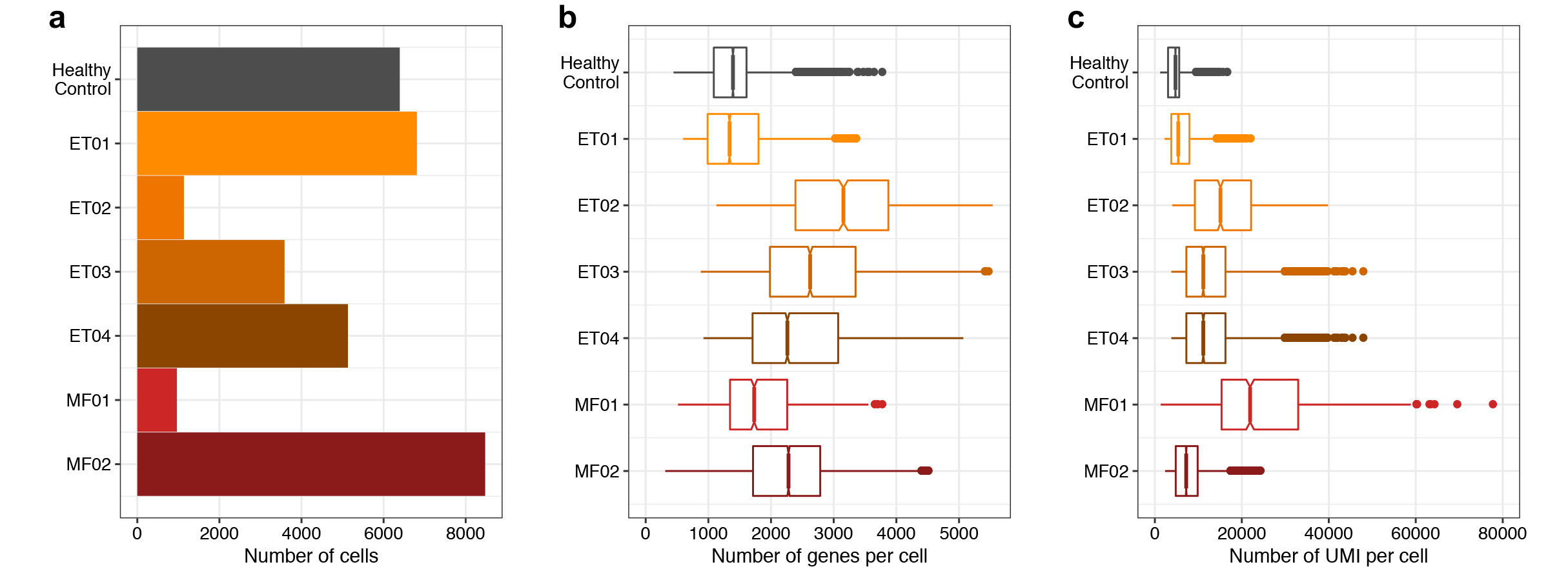
GoT does not compromise scRNA-seq data. (**a**) Number of cells, **(b)** number of genes per cell, and **(c)** number of UMIs per cell from published standard 10x data of healthy control CD34^+^ cells and 10x data of CD34^+^ cells from patient samples that underwent concurrent GoT.

**Supplementary Figure 7.**
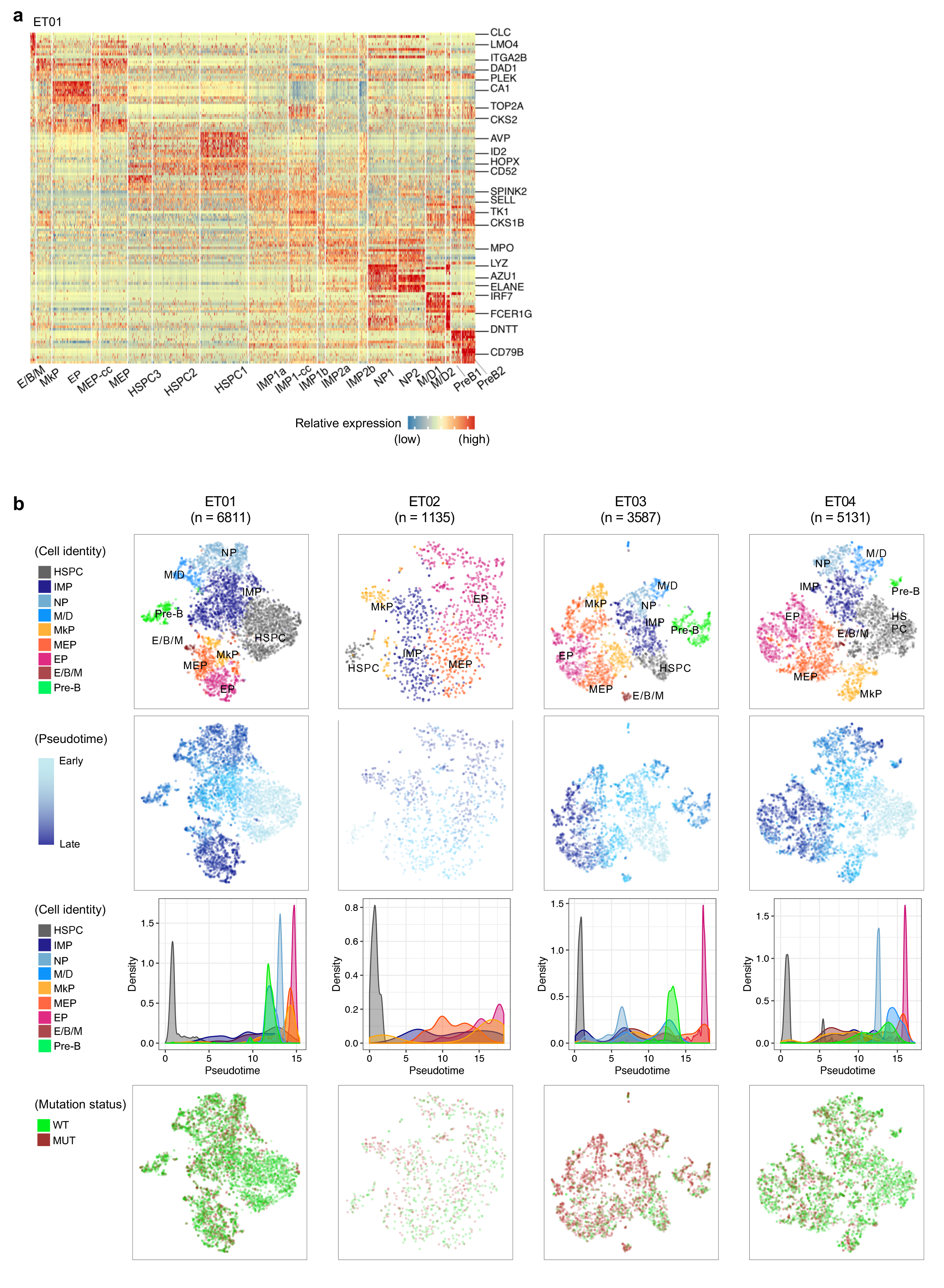
Combining progenitor cell identity and mutation status. **(a)** Example heatmap of differentially expressed genes between clusters, composed of lineage defining markers (Seurat package). **(b)** tSNE projections of ET patient samples with progenitor cluster assignment, pseudotime and mutation status. Density plots along the pseudotime points of each progenitor cell type. HSPC, hematopoietic stem progenitor cells; IMP, immature myeloid progenitors; NP, neutrophil progenitors; E/B/M, eosinophil, basophil, mast cell progenitors; MEP, megakaryocytic-erythroid progenitors; MkP, megakaryocytic progenitors; EP, erythroid progenitors; WT, wildtype; MUT, mutant.

**Supplementary Figure 8.**
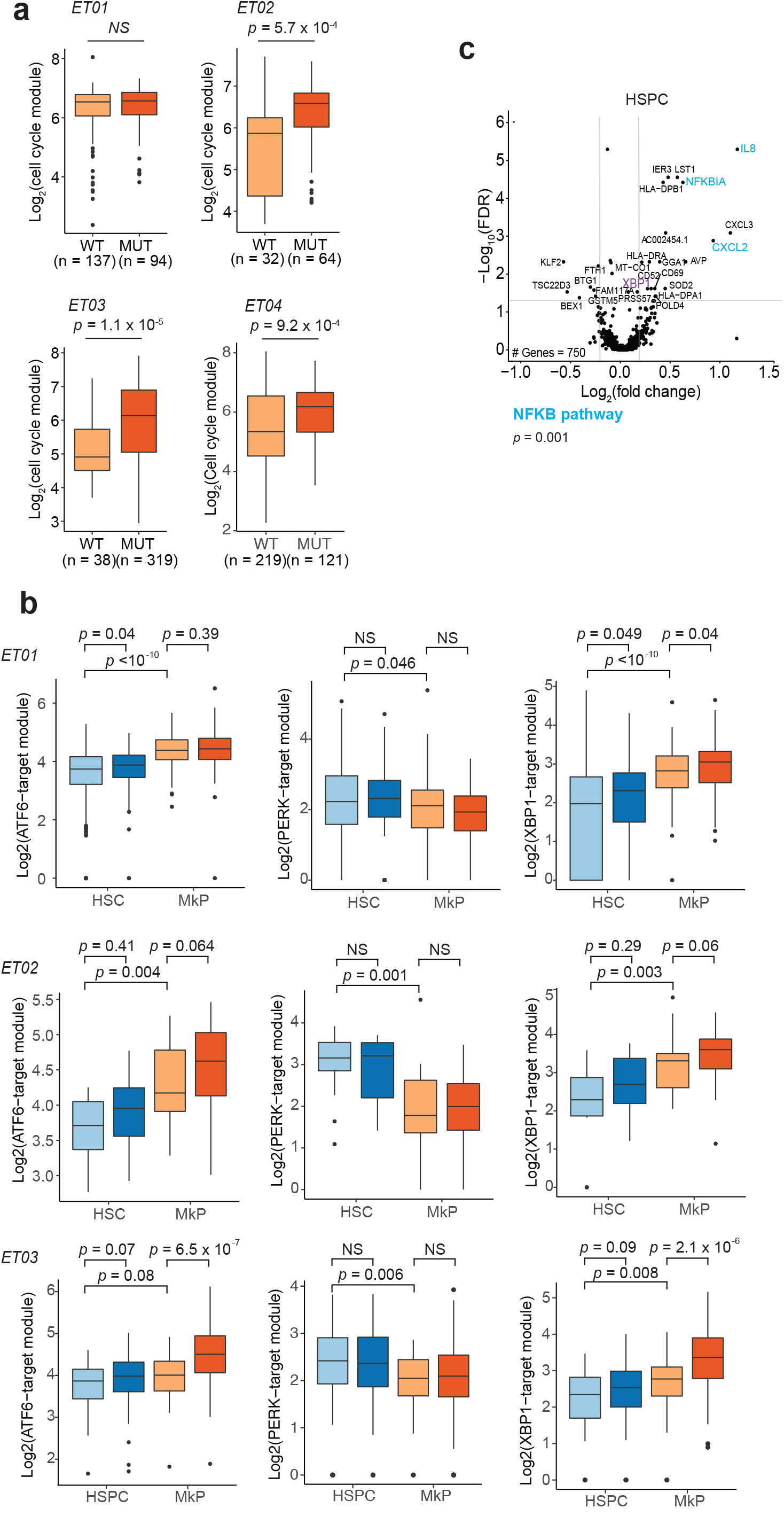
Gene expression analysis between *CALR* mutant and wildtype cells. **(a)** Cell cycle module score comparison between WT and MUT cells in the MkP clusters from ET patients (Wilcoxon rank-sum test). **(b)** Aggregate expression of ATF6-, PERK- and XBP1-target genes in the unfolded protein response in *CALR* WT and MUT HSPC and MkP cells. **(c)** Volcano plot of differentially expressed genes from patient sample ET01.

**Supplementary Figure 9.**
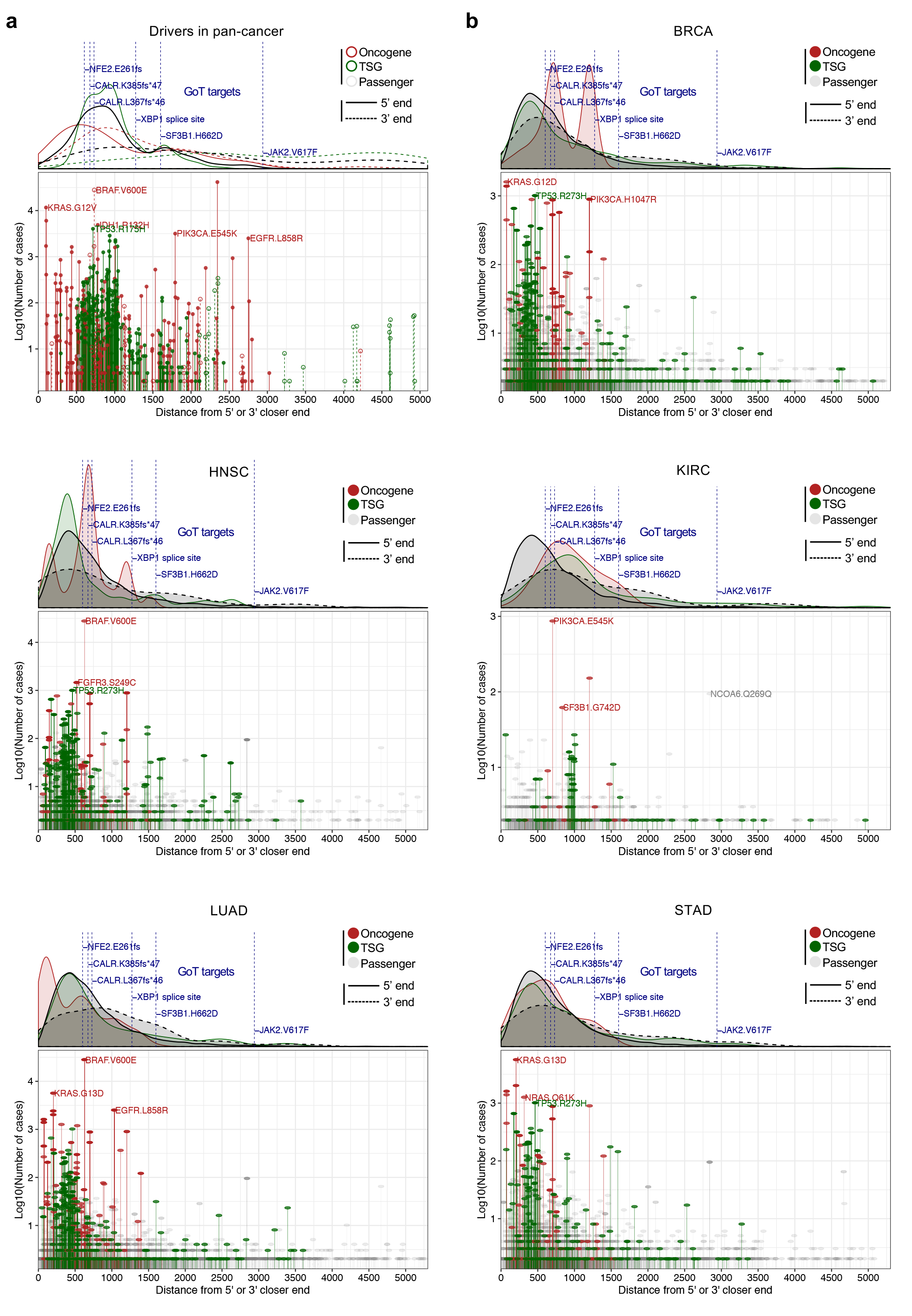
Distribution of mutation distance from 5’ or 3’ closer end of transcript. **(a)** Pan-cancer drivers, and **(b)** mutations identified in The Cancer Genome Atlas (TCGA) cohorts for the distribution of mutation distance from 5’ or 3’ closer end of the transcript, and their recurrent frequencies based on the number of times that they have been reported in Catalogue of Somatic Mutations in Cancer (COSMIC) database. Mutations are annotated as oncogenes, tumor suppressor genes (TSGs), or passengers (as defined in Vogelstein *et al*. 2013^71^ and Bailey *et al*. 2018^72^). Relative density of each subclass of mutations in the distance from 5’ or 3’ closer end is shown in the upper panel. GoT targets used in this study are annotated (in **(a)**, navy) to compare their distances relative to the references. BRCA, breast invasive carcinoma; HNSC, head and neck squamous cell carcinoma; KIRC, kidney renal clear cell carcinoma; LUAD, lung adenocarcinoma; STAD, stomach adenocarcinoma.

**Supplementary Figure 10.**
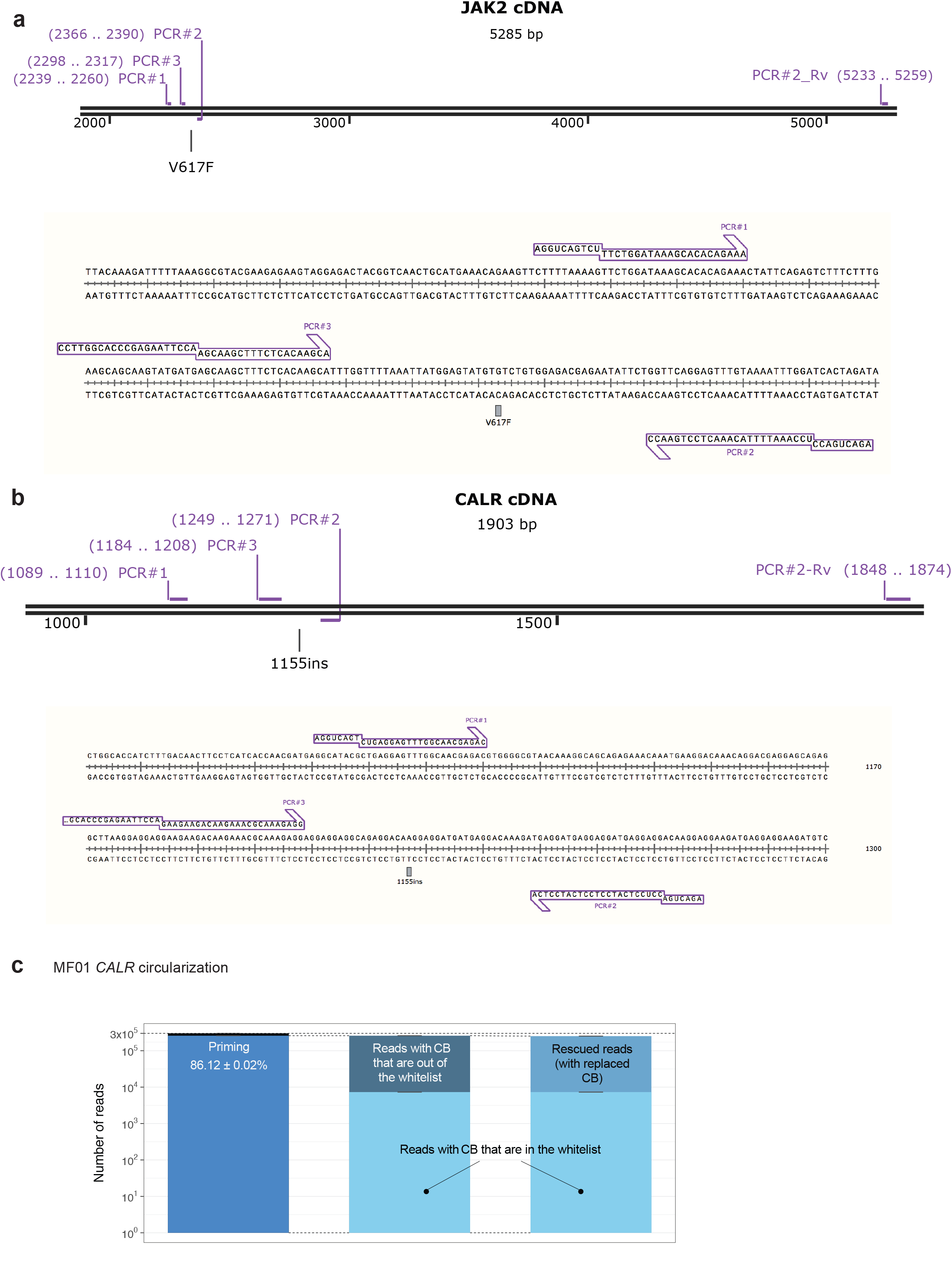
Circularization GoT captures targeted genotyping information. **(a)** Primer sequence positions for *JAK2* mutation in circularization GoT. **(b)** Primer sequence positions for type 2 *CALR* mutation in circularization GoT. **(c)** Targeted amplicon analysis result of circularization GoT from patient sample MF01. CB, cell barcode.

**Supplementary Figure 11.**
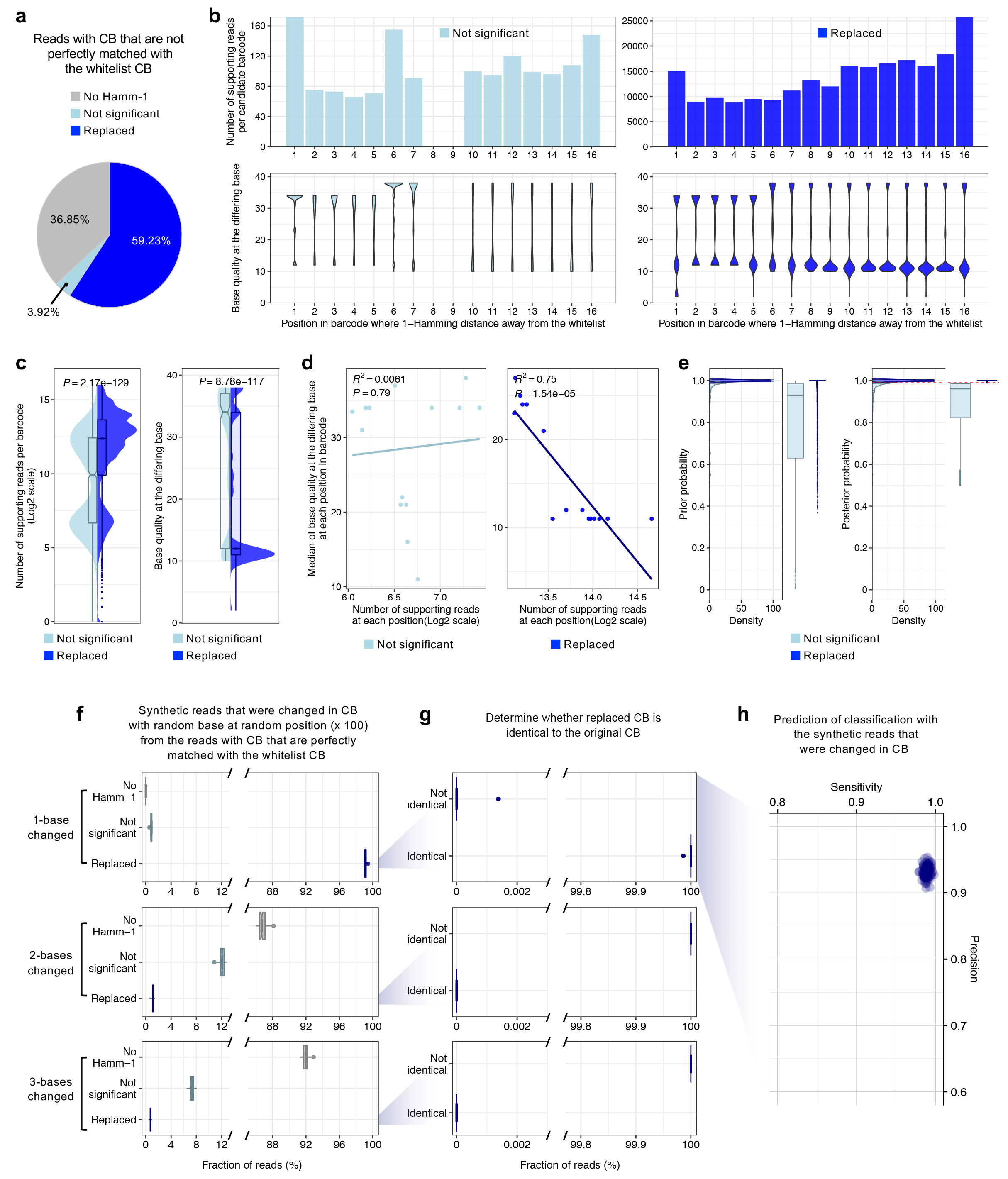
Evaluation of barcode replacement in GoT processing. **(a)** Fraction of reads with cell barcodes (CB) that are not perfectly matched with the whitelist CB from the species-mixing experiment. No Hamm-1, filtered reads with barcodes that have no replaceable candidates with 1-Hamming distance away from whitelisted barcodes; Not significant, filtered reads with barcodes that have candidates of 1-Hamming distance away from the whitelisted barcodes but no statistical significance (posterior probability < 0.99); Replaced, rescued reads with barcodes that have candidates of 1-Hamming distance away from the whitelisted barcodes with statistical significance (posterior probability ≥ 0.99). Number of supporting reads per candidate barcode and base quality **(b)** at each position in barcode where 1- Hamming distance away from the whitelist, and **(c)** across positions. Wilcoxon rank-sum tests were applied to compare two subclasses. **(d)** Correlation between the number of supporting reads per candidate barcode and median of base quality at each position in barcode where 1-Hamming distance away from the whitelist (two-tailed Pearson’s correlation). **(e)** Distribution of prior and posterior probabilities from two subclasses. A dashed red line represents the posterior probability cutoff (0.99). **(f-h)** To further evaluate the efficiency of barcode replacement, we generated synthetic reads that were changed in CB with random base at random position (× 100) from the reads with CB that are perfectly matched with the whitelist CB. **(f)** Fraction of reads with CB that are not perfectly matched with the whitelist CB from 100 iterations. Fractions of replaced reads were 99.1 ± 0.001% (median ± absolute deviation) in 1-base changed, 1.1 ± 0.002% in 2-bases changed, and 0.7 ± 0.001% in 3-bases changed simulations. **(g)** Determination of the fraction of reads with replaced CB that are identical or not to the original CB. In 1-base changed simulations, fraction of reads with replaced CB that were matched correctly to the original CB was 100.0 ± 0.0% (median ± absolute deviation). **(h)** Estimation of prediction power for classifying murine and human CB from 1-base changed simulations.

**Supplementary Table 1. Summary of patients’ clinical history and pathology.**

**Supplementary Table 2. Gene set module lists.**

**Supplementary Table 3. List of primers used in GoT.**

